# Gut bacterial tyrosine decarboxylases restrict the bioavailability of levodopa, the primary treatment in Parkinson’s disease

**DOI:** 10.1101/356246

**Authors:** Sebastiaan P. van Kessel, Alexandra K. Frye, Ahmed O. El-Gendy, Maria Castejon, Ali Keshavarzian, Gertjan van Dijk, Sahar El Aidy

## Abstract

Human gut bacteria play a critical role in the regulation of immune and metabolic systems, as well as in the function of the nervous system. The microbiota senses its environment and responds by releasing metabolites, some of which are key regulators of human health and disease. In this study, we identify and characterize gut-associated bacteria in their ability to decarboxylate L-DOPA (also known as Levodopa or L-3,4-dihydroxyphenylalanine) to dopamine via the tyrosine decarboxylases, which are mainly present in the class Bacilli. Although the bacterial tyrosine decarboxylases have a higher affinity for tyrosine compared to L-DOPA, this does not affect their ability to decarboxylate L-DOPA, nor does any inhibitor of the human decarboxylase. This study indicates that *in situ* bioavailability of L-DOPA is compromised by the gut bacterial tyrosine decarboxylase abundance in Parkinson’s patients. Finally, we show that the tyrosine decarboxylase abundance in the microbiota at the site of L-DOPA absorption, the proximal small intestine, significantly influences L-DOPA bioavailability in the plasma of rats. Our results highlight the role of microbial metabolism in drug bioavailability, and specifically, that small intestinal abundance of bacterial tyrosine decarboxylase can explain the highly variable L-DOPA dosage regimens required in the treatment of individual Parkinson’s patients.

**Highlights:**

- Small intestinal bacteria is able to convert L-DOPA to dopamine
- L-DOPA metabolism by gut bacteria reduce the bioavailability of L-DOPA in the body, thus is a significant explanatory factor of the highly variable L-DOPA dosage regimens required in the treatment of individual Parkinson’s patients.
- Inhibitors of the human DOPA decarboxylase are not potent inhibitors for bacterial tyrosine decarboxylases

## Introduction

The complex bacterial communities inhabiting the mammalian gut have a significant impact on the health of their host (Kahrstrom et al., 2016). Numerous reports indicate that intestinal microbiota, and in particular its metabolic products, have a crucial effect on various health and diseased states. Host immune system and brain development, metabolism, behavior, stress and pain response have all been reported to be associated with microbiota disturbances (El Aidy et al., 2012; Kelly et al., 2016; Mao et al., 2018; Pusceddu et al., 2015; Yano et al., 2015). In addition, it is becoming increasingly clear that gut microbiota can interfere with the modulation of drug pharmacokinetics and drug bioavailability (Enright et al., 2016; Niehues and Hensel, 2009).

Parkinson’s disease, the second most common neurodegenerative disorder, affecting 1% of the global population over the age of 60, has recently been correlated with alterations in microbial gut composition (Pereira et al., 2017; Sampson et al., 2016; Scheperjans et al., 2014). To date, L-DOPA is the most effective treatment available for Parkinson’s patients. In humans, peripheral L-DOPA metabolism involves DOPA decarboxylase, which converts L-DOPA to dopamine, thus preventing the passage of L-DOPA to its site of action in the brain, as dopamine cannot pass the blood-brain barrier (Pinder, 1970). For this reason, Parkinson’s patients are treated with a DOPA decarboxylase inhibitor (primarily carbidopa) in combination with L-DOPA to enhance the effectiveness of L-DOPA delivery to the brain (Deleu et al., 2002). Nonetheless, the pharmacokinetics of L-DOPA/carbidopa treatment varies significantly among patients, some are resistant to the treatment, others undergo fluctuating response towards the treatment over time, thus require increasing L-DOPA dosage regimen leading to increased severity of adverse effects like dyskinesia (Katzenschlager and Lees, 2002). What remains to be clarified is whether inter-individual variations in gut microbiota composition play a causative role in the variation of treatment efficacy. Several amino acid decarboxylases have been annotated in bacteria. Tyrosine decarboxylase genes (*tdc*) have especially been encoded in the genome of several bacterial species in the genera *Lactobacillus* and *Enterococcus* (Perez et al., 2015; Zhu et al., 2016). Though tyrosine decarboxylase (TDC) is named for its capacity to decarboxylate L-tyrosine into tyramine, it might also have the ability to decarboxylate L-DOPA to produce dopamine (Zhu et al., 2016) due to the high similarity of the chemical structures of these substrates. This implies that TDC activity of the gut microbiota might interfere with L-DOPA bioavailability, which could be of clinical significance in the L-DOPA treatment of Parkinson’s patients. The aim of the present study is to parse out the effect of L-DOPA metabolizing bacteria, particularly in the proximal small intestine, where L-DOPA is absorbed. Initially, we established TDC present in small intestinal bacteria efficiently converted L-DOPA to dopamine, confirming their capacity to modulate the *in situ* bioavailability of the primary drug used in the treatment of Parkinson’s patients. We show that higher relative abundance of bacterial *tdc* gene in fecal samples of Parkinson’s patients positively correlates with higher daily L-DOPA dosage requirement and duration of disease. We further confirm our findings in rats orally administered a mixture of L-DOPA/carbidopa, illustrating that L-DOPA levels in plasma negatively correlate with the abundance of bacterial *tdc* gene in the proximal small intestine.

## Results

### Proximal small intestinal bacteria of rodents convert L-DOPA to dopamine

To determine whether proximal small intestinal microbiota maintain the ability to metabolize L-DOPA, luminal samples from the jejunum of wild-type Groningen rats were incubated *in vitro* with L-DOPA and analyzed by High-Performance Liquid Chromatography with Electrochemical Detection (HPLC-ED). The chromatograms revealed that L-DOPA decarboxylation (**Figure 1A**) coincides with the conversion of tyrosine to tyramine (**Figure 1B-E**). In addition, no other metabolites derived from L-DOPA were detected. To support the *ex vivo* experiment results, the uptake of L-DOPA was quantified in plasma samples from specific pathogen free and germ-free female C57 BL/6J mice after oral gavage with L-DOPA. HPLC-ED analysis revealed higher levels of L-DOPA and its metabolites dopamine and DOPAC (3,4-Dihydroxyphenylacetic acid) in plasma samples of germ-free mice compared to their conventional counterparts (**Figure S1**). Taken together, the results suggest that TDC is involved in L-DOPA metabolism by gut bacteria, which may, in turn, interfere with L-DOPA uptake in the proximal small intestine.

**Figure 1.**
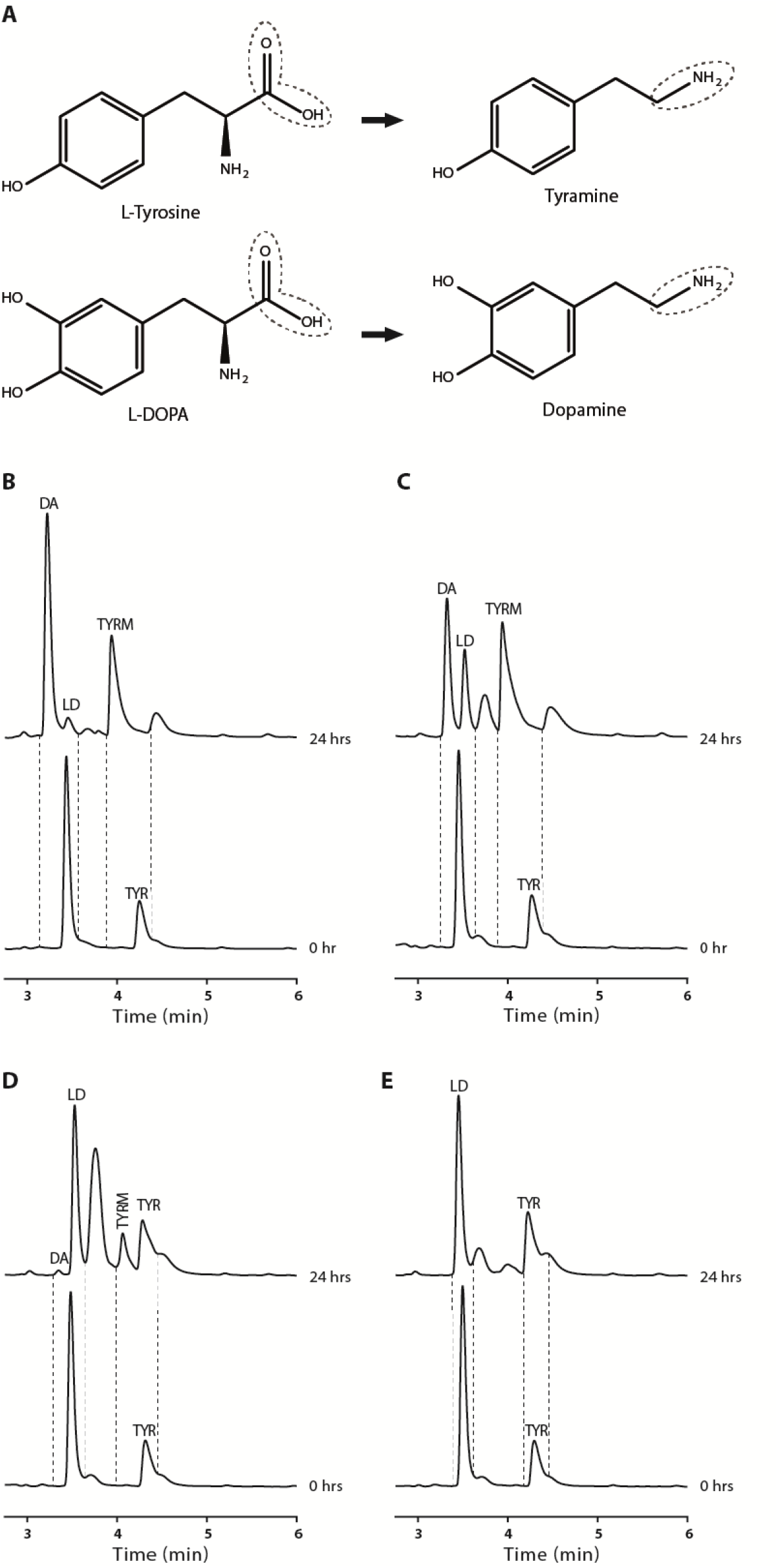
Bacteria in jejunal content decarboxylate L-DOPA to dopamine coinciding with their production of tyramine *ex vivo*. (**A**) Decarboxylation reaction for tyrosine and L-DOPA. (**B-E**) Jejunal extracts from 4 different rats. (**B, C**) Bacterial conversion of tyrosine (TYR) to tyramine (TYRM) and 1 mM of supplemented L-DOPA (LD) to dopamine (DA), during 24 hrs of incubation. (**D**) Detection of low amounts of dopamine production when tyrosine was still abundant and tyramine production was relatively low. (**E**) No tyrosine was converted to tyramine; accordingly, no L-DOPA was converted to dopamine.

### Bacterial tyrosine decarboxylase is responsible for L-DOPA decarboxylation

The coinciding tyrosine and L-DOPA decarboxylation observed in the luminal content of jejunum was the basis of our hypothesis that TDC is the enzyme involved in both conversions. Species of the genera *Lactobacillus* and *Enterococcus* have been reported to harbor this enzyme (Perez et al., 2015; Zhang and Ni, 2014). To investigate whether genomes of these genera indeed represent the main TDC encoding bacterial groups of the small intestine microbiota, the TDC protein (EOT87933) from the laboratory strain *Enterococcus faecalis* v583 was used as a query to search the US National Institutes of Health Human Microbiome Project (HMP) protein database, to identify whether the genome of other (small intestinal) gut bacteria also encode *tdc*. This analysis exclusively identified TDC proteins in species belonging to the bacilli class, including more than 50 *Enterococcus* strains (mainly *E. faecium* and *E. faecalis*) and several *Lactobacillus* and *Staphylococcus* species (**Figure S2A**).

Next, we aligned the genome of *E. faecalis* v583 with two gut bacterial isolates, *E. faecium* W54, and *L. brevis* W63, illustrating the conservation of the *tdc*-operon among these species (**Figure 2A**). Intriguingly, analysis of *E. faecium* genomes revealed that this species encodes a second, paralogous *tdc* gene (^P^TDC_EFM_) that did not align with the conserved *tdc*-operon and was absent from the other species (**Figure 2A, Figure S2A, Data file S1**).

**Figure 2.**
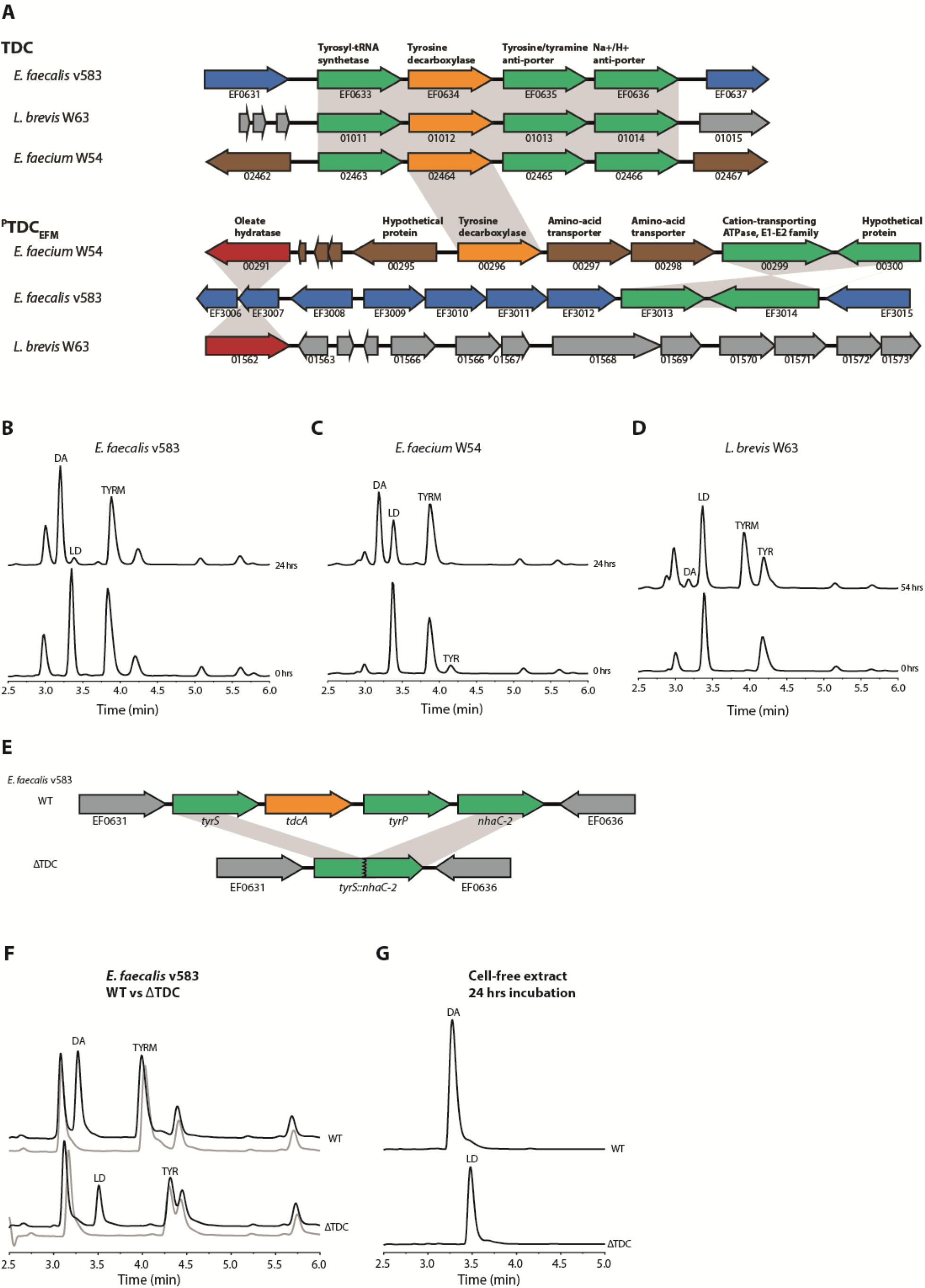
Gut bacteria harboring tyrosine decarboxylases are responsible for L-DOPA decarboxylation. (**A**) Aligned genomes of *E. faecium, E. faecalis*, and *L. brevis.* The conserved *tdc*-operon is depicted with *tdc*-gene in orange. Overnight cultures of (**B**) *E. faecalis* v583, (**C**) *E. faecium* W54, and (**D**) *L. brevis* W63 incubated anaerobically at 37 °C with 100 µM of L-DOPA (LD). (**E**) Genomic scheme showing the differences between EFS^WT^ (this study) and EFS^ΔTDC^(Perez et al., 2015). (**F**) Overnight cultures of EFS^WT^ and EFS^ΔTDC^ incubated anaerobically at 37 °C with 100 uM L-DOPA (black line) compared to control (grey line) where no L-DOPA was added. (**G**) Comparison of enzyme activity of EFS^WT^ and EFS^ΔTDC^ cell-free protein extracts after O/N incubation with 1 mM L-DOPA at 37 °C. Curves represent one example of 3 biological replicates.

To support our *in silico* data, a comprehensive screening of *E. faecalis* v583, *E. faecium* W54, and *L. brevis* W63 and 77 additional clinical and human isolates of *Enterococcus*, including clinical isolates and strains from healthy subjects, was performed. All enterococcal isolates and *L. brevis* were able to convert tyrosine and L-DOPA into tyramine and dopamine, respectively (**Figure 2B-D, Table S1**). Notably, our HPLC-ED analysis revealed considerable variability among the tested strains with regard to their efficiency to decarboxylate L-DOPA. *E. faecium* and *E. faecalis* were drastically more efficient at converting L-DOPA to dopamine, compared to *L. brevis*. Growing *L. brevis* in different growth media did not change the L-DOPA decarboxylation efficacy (**Figure S2B, C**). To eliminate the possibility that other bacterial amino acid decarboxylases are involved in L-DOPA conversion observed in the jejunal content we expanded our screening to include live bacterial species harboring PLP-dependent amino acid decarboxylases previously identified by Williams et al (Williams et al., 2014). None of the tested bacterial strains encoding different amino acid decarboxylases could decarboxylate L-DOPA (**Figure S2D-G, Table S2**).

To verify that the *tdc* gene is solely responsible for L-DOPA decarboxylation in *Enterococcus*, wild type *E. faecalis* v583 (EFS^WT^) was compared with a mutant strain (EFS^ΔTDC^) in which both the *tdc* gene (*tdcA*) and tyrosine transporter (*tyrP*) were deleted (*14*), (**Figure 2E**). Overnight incubation of EFS^WT^ and EFS^ΔTDC^ bacterial cells with L-DOPA resulted in production of dopamine in the supernatant of EFS^WT^ but not EFS^ΔTDC^ (**Figure 2F**), confirming the pivotal role of these genes in this conversion. To rule out that deletion of *tyrP* alone could explain the observed result by an impaired L-DOPA import, cell-free protein extracts were incubated with 1 mM L-DOPA overnight at 37°C. While EFS^WT^ cell-free protein extract converted all supplied L-DOPA into dopamine, no dopamine production was observed in the EFS^ΔTDC^ cell-free protein extracts (**Figure 2G**). Collectively, results show that TDC is encoded on genomes of gut bacterial species known to dominate the proximal small intestine and that this enzyme is exclusively responsible for converting L-DOPA to dopamine by these bacteria, although the efficiency of that conversion displays considerable species-dependent variability.

### High levels of tyrosine do not prevent bacterial decarboxylation of L-DOPA

To test whether the availability of the primary substrate for bacterial tyrosine decarboxylases (i.e., tyrosine) could inhibit the uptake and decarboxylation of L-DOPA, the growth, metabolites, and pH that was previously shown to affect the expression of *tdc* (Perez et al., 2015), of *E. faecium* W54 and *E. faecalis* v583 were analyzed over time. 100 µM L-DOPA was added to the bacterial cultures, whereas approximately 500 µM tyrosine was present in the growth media. Remarkably, L-DOPA and tyrosine were converted simultaneously, even in the presence of these excess levels of tyrosine (1:5 L-DOPA to tyrosine), albeit at a slower conversion rate for L-DOPA (**Figure 3A-B**). Notably, the decarboxylation reaction appeared operational throughout the exponential phase of growth for *E. faecalis*, whereas it is only observed in *E. faecium* when this bacterium entered the stationary phase of growth, suggesting differential regulation of the *tdc* expression in these species.

**Figure 3.**
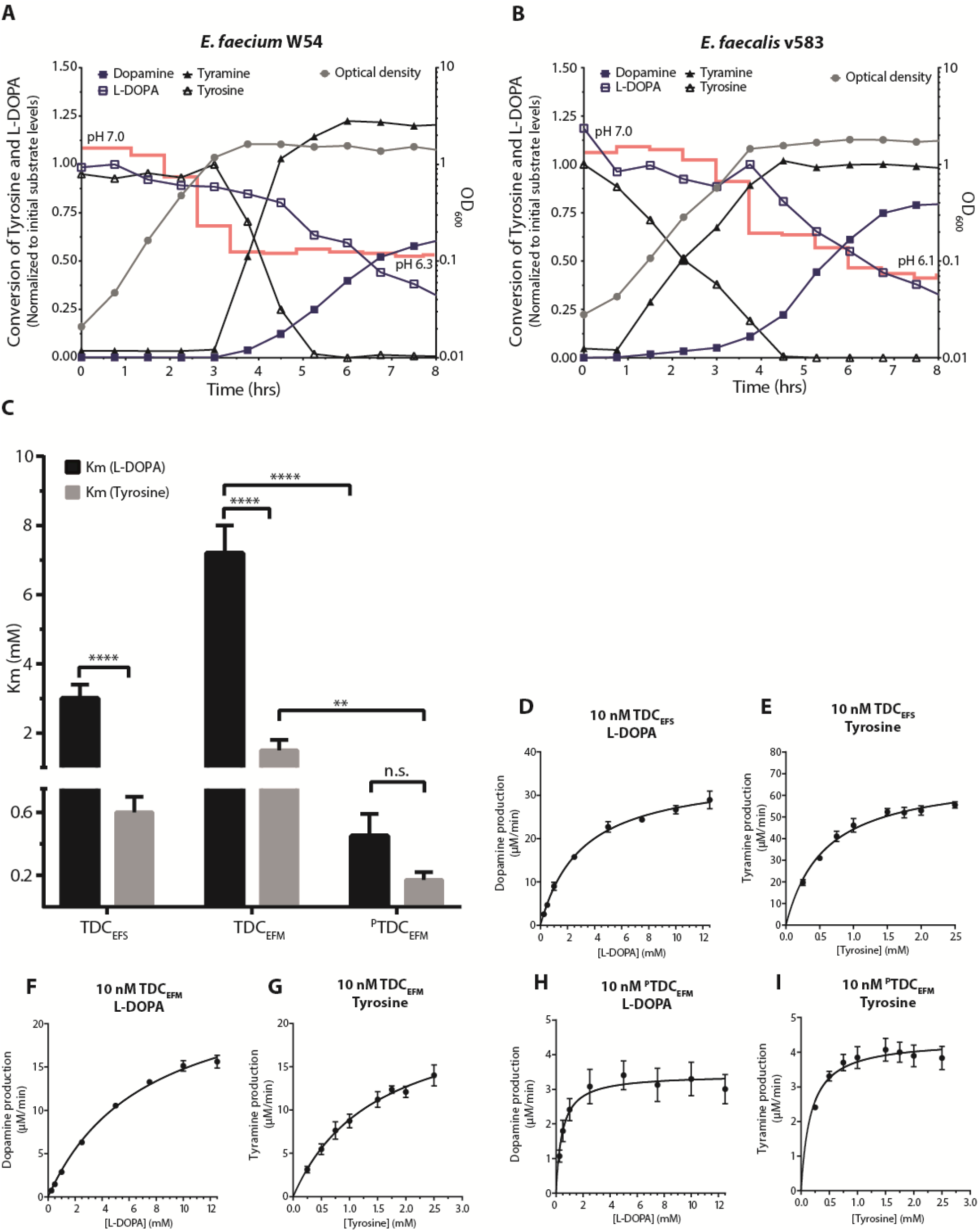
Enterococci decarboxylate L-DOPA in presence of tyrosine despite higher affinity for tyrosine *in vitro*. Growth curve (grey circle, right Y-axis) of *E. faecium* W54 (**A**) and *E. faecalis* (**B**) plotted together with L-DOPA (open square), dopamine (closed square), tyrosine (open triangle), and tyramine (closed triangle) levels (left Y-axis). Concentrations of product and substrate were normalized to the initial levels of the corresponding substrate (100 µM supplemented L-DOPA and ∼500 µM tyrosine present in the medium). pH of the culture is indicated over time as a red line. (**C**) Substrate affinity (Km) for L-DOPA and tyrosine for purified tyrosine decarboxylases from *E. faecalis* v583 (TDC_EFS_), *E. faecium* W54 (TDC_EFM_, PTDC_EFM_). (**D-I**) Michaelis-Menten kinetic curves for L-DOPA and tyrosine as substrate for TDC_EFS_ (**D,E**), TDC_EFM_ (**F,G**), and ^P^TDC_EFM_ (**H,I**). Reactions were performed in triplicate using L-DOPA concentrations ranging from 0.5-12.5 mM and tyrosine concentrations ranging from 0.25-2.5 mM. The enzyme kinetic parameters were calculated using nonlinear Michaelis-Menten regression model. Error bars represent the SEM and significance was tested using 2-way-Anova, Fisher LSD test, (*=p<0.02 **=p<0.01 ****<0.0001).

To further characterize the substrate specificity and kinetic parameters of the bacterial tyrosine decarboxylases, *tdc* genes from *E. faecalis* v583 (TDC_EFS_) and *E. faecium* W54 (TDC_EFM_ and ^P^TDC_EFM_) were expressed in *Escherichia coli* BL21 (DE3) and then purified. Michaelis-Menten kinetics indicated each of the studied enzymes had a significantly higher affinity (K_m_) (**Figure 3C-I**) and catalytic efficiency (K_cat_/K_m_) for tyrosine than for L-DOPA (**Table 1**). Despite the differential substrate affinity, our findings illustrate that high levels of tyrosine do not prevent the decarboxylation of L-DOPA in batch culture.

**Table 1.** Michaelis-Menten kinetic parameters. Enzyme kinetic parameters were determined by Michaelis-Menten nonlinear regression model for L-DOPA and Tyrosine as substrates. ± indicates the standard error.

| **L-DOPA** | pH 5.0 | pH 5.0 | pH 4.5 | pH 7.4 |
| --- | --- | --- | --- | --- |
|  | **TDC<sub>EFS</sub>** | **TDC<sub>EFM</sub>** | **<sup>P</sup>TDC<sub>EFM</sub>** | **DDC** |
| [E] (nM) | 10.00 | 10.00 | 10.00 | 10.00 |
| K<sub>m</sub> (mM) | 3±0.4 | 7.2±0.8 | 0.4±0.1 | 0.1±0.01 |
| V<sub>max</sub> (μM/min) | 35.3±1.4 | 25.5±1.3 | 3.4±0.2 | 1.4±0.03 |
| K<sub>cat</sub> (min<sup>-1</sup>) | 3531±137 | 2549±133 | 342.4±21 | 136.9±3 |
| K<sub>cat</sub>/K<sub>m</sub> (min<sup>-1</sup>/mM<sup>-1</sup>) | 1160 | 352 | 764 | 1567 |
| R<sup>2</sup> | 0.978 | 0.990 | 0.621 | 0.962 |

| **Tyrosine** | pH 5.0 | pH 5.0 | pH 4.5 |
| --- | --- | --- | --- |
|  | **TDC<sub>EFS</sub>** | **TDC<sub>EFM</sub>** | **<sup>P</sup>TDC<sub>EFM</sub>** |
| [E] (nM) | 10.00 | 10.00 | 10.00 |
| K<sub>m</sub> (mM) | 0.6±0.1 | 1.5±0.3 | 0.2±0.05 |
| V<sub>max</sub> (μM/min) | 69.6±2.9 | 22±2.5 | 4.4±0.2 |
| K<sub>cat</sub> (min<sup>-1</sup>) | 6963±288 | 2204±247 | 435.6±19.2 |
| K<sub>cat</sub>/K<sub>m</sub> (min<sup>-1</sup>/mM<sup>-1</sup>) | 12216 | 1493 | 2558 |
| R<sup>2</sup> | 0.928 | 0.902 | 0.589 |

### Carbidopa is a potent inhibitor of the human decarboxylase but not of bacterial decarboxylases

To assess the extent to which human DOPA decarboxylase inhibitors could affect bacterial decarboxylases, three human DOPA decarboxylase inhibitors (carbidopa, benserazide, and methyldopa) were tested on purified bacterial TDCs and on the corresponding bacterial batch cultures. Comparison of the inhibitory constants (K_i_^TDC^/K_i_^DDC^) demonstrates carbidopa to be a 1.4-1.9 x 10^4^ times more potent inhibitor of human DOPA decarboxylase than bacterial TDCs (**Figure 4A, S3; Table S3**). This is best illustrated by the observation that L-DOPA conversion by *E. faecium* W54 and *E. faecalis* v583 batch cultures (OD_600_= ∼2.0) was unaffected by co-incubation with carbidopa (equimolar or 4-fold carbidopa relative to L-DOPA) (**Figure 4B-C**, **S4A**). Analogously, benserazide and methyldopa did not inhibit the L-DOPA decarboxylation activity in *E. faecalis* or *E. faecium* (**Figure S4B-C**).

**Figure 4.**
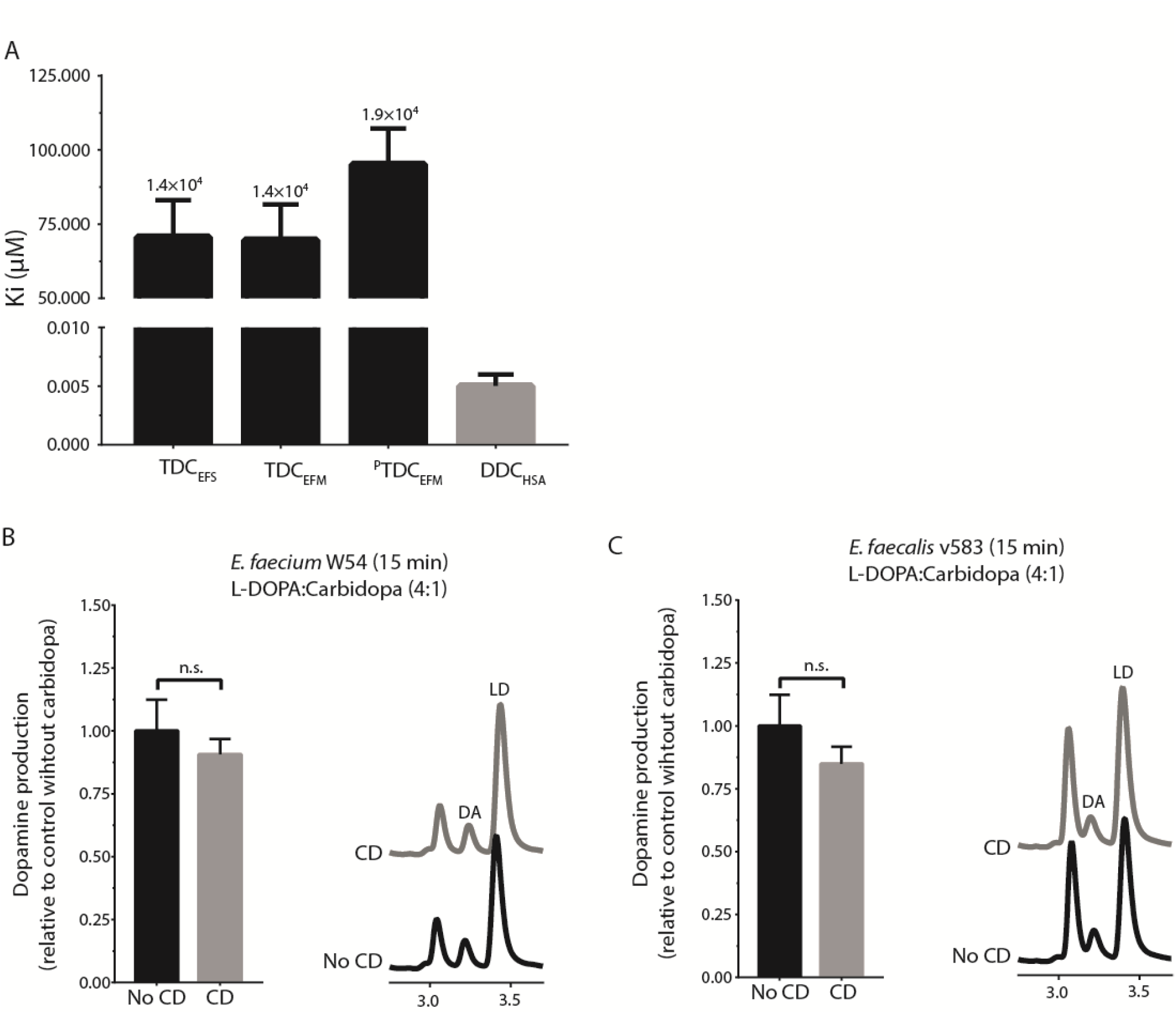
Human DOPA decarboxylase inhibitor, carbidopa, does not inhibit bacterial tyrosine decarboxylases. (**A**) Inhibitory constants (Ki) of bacterial decarboxylases (black) and human DOPA decarboxylase (grey), with fold-difference between bacterial and human decarboxylase displayed on top of the bars. Quantitative comparison of dopamine (DA) production by *E. faecium* W54, (**B**) and *E. faecalis* v583, (**C**) at stationary phase after 15 min, with representative HPLC-ED curve. Bacterial cultures (n=3) were incubated with 100 µM L-DOPA (LD) or a 4:1 mixture (in weight) of L-DOPA and carbidopa (CD) (100 µM L-DOPA and 21.7 µM carbidopa). Longer incubations resulted in complete conversion of L-DOPA to dopamine also in the presence of carbidopa (Data not shown). Error bars represent SEM (**A**) or SD (**B,C**) and significance was tested using a parametric unpaired T-test.

These findings demonstrate the commonly applied inhibitors of human DOPA decarboxylase in L-DOPA combination therapy do not inhibit bacterial TDC dependent L-DOPA conversion, implying L-DOPA/carbidopa combination therapy for Parkinson’s patients would not affect the bioavailability of L-DOPA *in situ* by intestinal bacteria.

### L-DOPA dosage regimen correlate with tyrosine decarboxylase gene abundance in Parkinson’s patients

To determine whether the considerable variation in L-DOPA dosage required for individual Parkinson’s patients could be attributed to the abundance of *tdc* genes in the gut microbiota, fecal samples were collected from male and female Parkinson’s patients (**Table S4**) on different doses of L-DOPA/carbidopa treatment (ranging from 300 up to 1100 mg L-DOPA per day). *tdc* gene-specific primers were used to quantify its relative abundance within the gut microbiota by qPCR (**Figure S5**). Remarkably, Pearson *r* correlation analyses showed a strong positive correlation (*r* = 0.70, R^2^ = 0.49, *p value* = 0.024) between bacterial *tdc* gene relative abundance and L-DOPA treatment dose (**Figure 5A**) as well as with the duration of disease (**Fig. 5B**). At this stage, it is unclear whether the relative abundance of *tdc* genes in fecal samples reflects its abundance in the small intestinal microbiota. This is of particular importance because L-DOPA is absorbed in the proximal small intestine, and reduction in its bioavailability by bacterial TDC activity in the context of Parkinson’s patients’ medication regimens would only be relevant in that intestinal region. Still, the selective prevalence of *tdc* encoding genes in the genomes of signature microbes of the small intestine microbiota supports the notion that the results obtained from fecal samples are a valid representation of *tdc* gene abundance in the small intestinal microbiota. Moreover, the significant correlation of the relative *tdc* abundance in the fecal microbiota and the required L-DOPA dosage as well as disease duration strongly supports a role for bacterial TDC in L-DOPA bioavailability.

**Figure 5.**
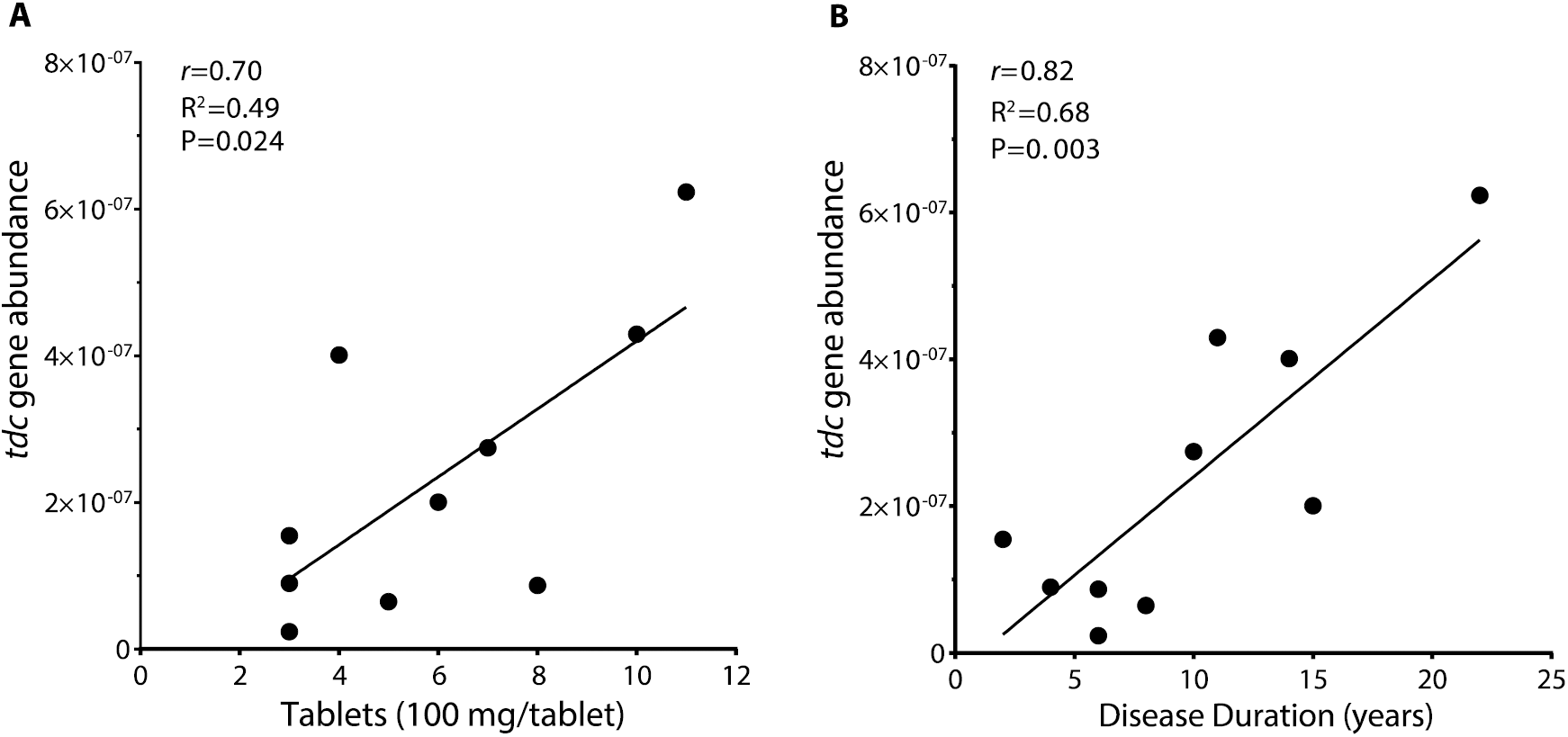
Tyrosine decarboxylase gene abundance correlates with daily L-DOPA dose and disease duration in fecal samples of Parkinson’s patients. (A) Scatter plot of *tdc* gene abundance measured by qPCR in fecal samples of Parkinson’s patients (n=10) versus daily L-DOPA dosage fitted with linear regression model. (B) Scatter plot of *tdc* gene abundance from the same samples versus disease duration fitted with a linear regression model. Pearson’s r analysis was used to determine significant correlations between tyrosine decarboxylase -gene abundance and dosage (r=0.70, R2=0.49, P value=0.024) or disease duration (r = 0.82, R2 = 0.68, P value = 0.003).

### Tyrosine decarboxylase gene abundance in small intestine correlates with L-DOPA bioavailability in rats

To further consolidate the concept that *tdc* gene abundance in proximal small intestinal microbiota affects peripheral levels of L-DOPA in blood and dopamine/L-DOPA ratio in the jejunal luminal content, male wild-type Groningen rats (n=25) rats were orally administered 15 mg L-DOPA/3.75 mg carbidopa per kg of body weight and sacrificed after 15 minutes (point of maximal L-DOPA bioavailability in rats (Bredberg et al., 1994)). Plasma levels of L-DOPA and its metabolites dopamine and DOPAC were measured by HPLC-ED, while relative abundance of the *tdc* gene within the small intestinal microbiota was quantified by gene-specific qPCR (**Figure S5**). Strikingly, Pearson *r* correlation analyses showed that the ratio between dopamine and L-DOPA levels in the proximal jejunal content positively correlated with *tdc* gene abundance (*r*= 0.78, R^2^= 0.61, *P* value = 0.0001) (**Figure 6A**), whereas the absolute L-DOPA concentration in the proximal jejunal content was negatively correlated with the abundance of the *tdc* gene (*r*= −0.68, R^2^= 0.46, *P* value = 0.0021) (**Figure 6B**). Moreover, plasma levels of L-DOPA displayed a strong negative correlation (r = −0.66, R^2^ = 0.434, *P* value = 0.004) with the relative abundance of the *tdc* gene (**Figure 6C**). Findings indicate that L-DOPA uptake by the host is compromised by higher abundance of gut bacteria encoding for *tdc* genes in the upper region of the small intestine.

**Figure 6.**
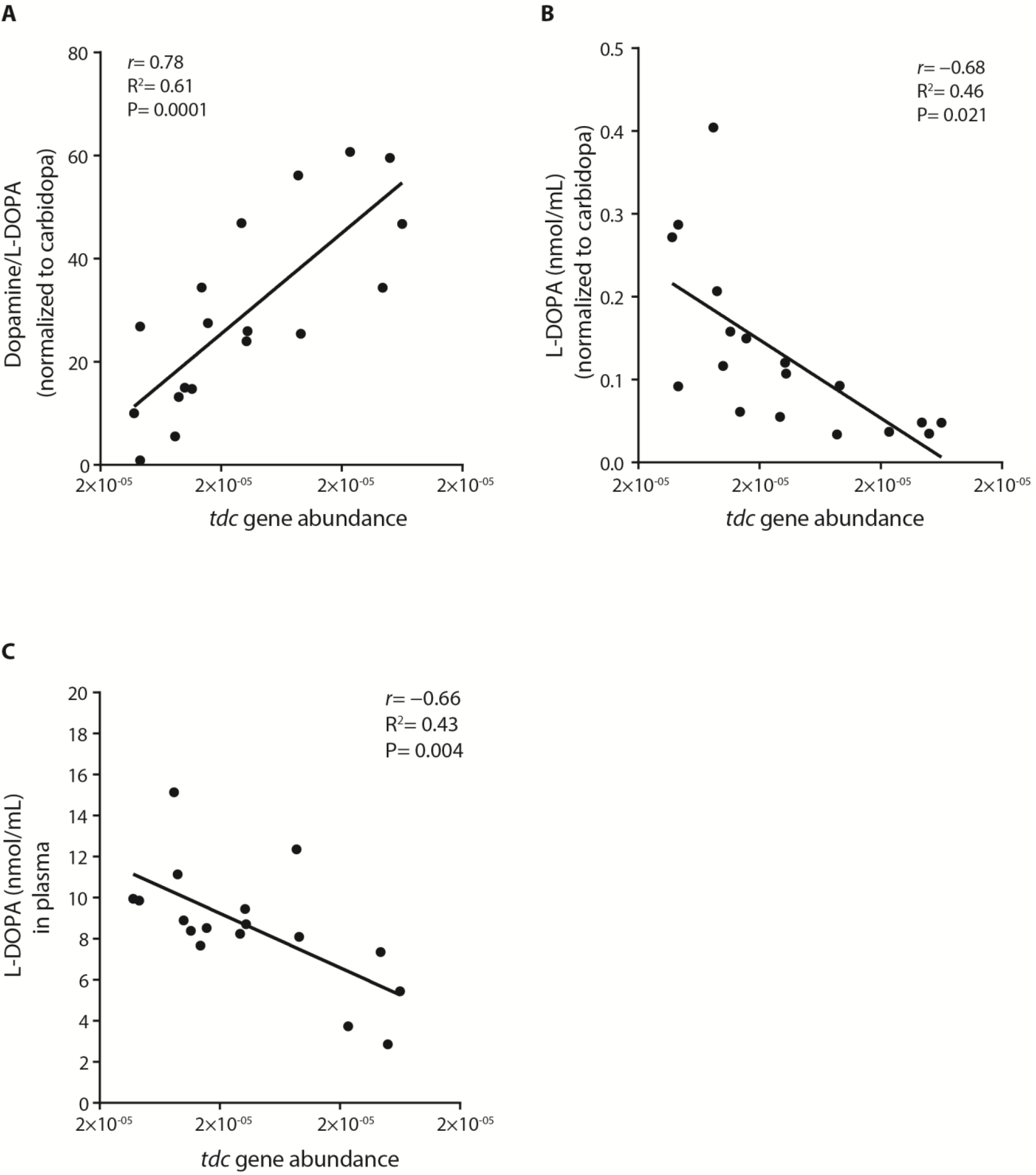
Tyrosine decarboxylase gene abundance correlates with L-DOPA bioavailability in rats. Scatter plot of *tdc* gene abundance measured by qPCR in jejunal content of wild-type Groningen rats (n=18) orally supplied with a L-DOPA/carbidopa mixture (4:1) versus (**A**) the dopamine/L-DOPA levels in the jejunal content, (**B**) the L-DOPA levels in the jejunal content, (**C**) or the L-DOPA levels in the plasma, fitted with a linear regression model. Pearson’s *r* correlation was used to determine significant correlations between TDC-gene abundance and jejunal dopamine levels (r = 0.78, R^2^ = 0.61, *P* value = 0.0001), jejunal L-DOPA levels (r = −0.68, R^2^ = 0.46 *P* value = 0.021), or plasma L-DOPA levels (r = −0.66, R^2^ = 0.43, *P* value = 0.004). No L-DOPA, dopamine, or DOPAC were detected in the control group (n=5).

## Discussion

Our observation of small intestinal microbiota able to convert L-DOPA to dopamine (**Figure 1**) was the basis of investigating the role of L-DOPA metabolizing bacteria in the context of the disparity in effective L-DOPA treatment between Parkinson’s patients (**Figure 5**) for which an appropriate explanation is lacking (Tomlinson et al., 2010). This study identifies a significant factor to consider in both the evaluation and treatment efficacy of L-DOPA/carbidopa pharmacokinetics and pharmacodynamics. Our primary outcome is that L-DOPA decarboxylation by gut bacteria, particularly if found in higher abundances in the proximal small intestine, would drastically reduce the bioavailability of L-DOPA in the body, and thereby contribute to the observed higher dosages required in some patients. Previously, reduced L-DOPA availability has been associated with *Helicobacter pylori* positive Parkinson’s patients, which was explained by the observation that *H. pylori* could bind L-DOPA *in vitro* via surface adhesins (Niehues and Hensel, 2009). However, this explanation is valid only for a small population of the Parkinson’s patients, who suffer from stomach ulcers and thus have high abundance of *H. pylori.*

Parkinson’s patients also often suffer from impaired small intestinal motility (Pellegrini et al., 2015), and are frequently administered proton pump inhibitors (PPIs) (Richter, 2007), leading to small intestinal bacterial overgrowth (Pfeiffer, 2013; Tan et al., 2014). Members of bacilli, including the genera *Enterococcus* and *Lactobacillus*, were previously identified as the predominant residents of the small intestine (El Aidy et al., 2014; Zoetendal et al., 2012), and in particular, *Enterococcus* has been reported to dominate in proton pump inhibitors’ induced small intestinal bacterial overgrowth (Freedberg et al., 2015). These factors contribute to a higher abundance of bacteria harboring the *tdc* gene in the small intestine, which would reduce the bioavailability of L-DOPA. Moreover, the correspondingly rising levels of dopamine may further aggravate the reduced gut motility as previously shown (Valenzuela and Dooley, 1984), thereby enhancing a state of small intestinal bacterial overgrowth, and perpetuating a vicious circle leading to higher L-DOPA dosage requirement for effective treatment of individual Parkinson’s patients. Alternatively, prolonged L-DOPA administration may influence tyrosine decarboxylase-gene abundance by selectively stimulating TDC harboring bacteria in the gut. In fact, it has been shown that the fitness of *E. faecalis* v583 in low pH depends on the *tdc*-operon (Perez et al., 2015), indicating long-term exposure to L-DOPA could contribute to selection for overgrowth of *tdc* encoding bacteria *in vivo* as supported by the positive correlation with with disease duration and *tdc* gene abundance observed in human fecal samples (**Figure 5**). This would explain the fluctuating response to L-DOPA during prolonged disease treatment, and the consequent increased L-DOPA dosage regimen leading to increased severity of its adverse effects such as dyskinesia (Kempster et al., 2007).

While our further investigation into the kinetics of both bacterial and human decarboxylases support the effectiveness of carbidopa to inhibit the human DOPA decarboxylase, it also shows that the same drug fails to inhibit L-DOPA decarboxylation by bacterial TDC, probably due to the presence of an extra hydroxyl group on the benzene ring of Carbidopa (**Figure 4, S4**). This suggests a better equilibration of L-DOPA treatment between patients could potentially be achieved by co-administration of an effective TDC inhibitor that targets both human and bacterial decarboxylases. Alternatively, we are currently evaluating regulation of *tdc* gene expression to help avoid the need for high L-DOPA dosing, thus minimizing its adverse side effects.

Notably, a few enterococcal strains that harbor the *tdc* gene are marked as probiotics. The use of such strains as dietary supplements should be recognized in case of Parkinson’s patients. More generally, our data support the increasing interest in the impact that gut microbiota metabolism may have on medical treatment and diet.

Collectively, our data show that L-DOPA conversion by bacterial TDC in the small intestine should be considered as a significant explanatory factor for the highly variable L-DOPA dosage regimens required in the treatment of individual Parkinson’s patients. These bacteria or their encoded *tdc* gene may potentially serve as a predictive biomarker for patient stratification to predict and consequently adjust the effective L-DOPA dosage regimens for Parkinson’s patients on an individual basis. Such biomarker potential is supported by the significant and robust (*r*=0.70) correlation observed between the relative abundance of bacterial *tdc* genes in fecal samples of patients and number of L-DOPA tablets required to treat individual Parkinson’s patients (**Figure 5**).

A limitation of the present study is the relatively small number of patient fecal samples analyzed, and the lack of evidence as to whether high abundance of *tdc in vivo* is a cause or consequence of higher L-DOPA dosage requirement during prolonged disease treatment. To overcome these limitations, a large longitudinal cohort of newly diagnosed Parkinson’s patients should be designed and followed over long periods of time, with *tdc* gene abundance employed as a personal microbiota-based biomarker to predict individual L-DOPA dosage requirement.

## Acknowledgements

We thank Dr. Saskia van Hemert and Dr. Coline Gerritsen of Winclove Probiotics, Amsterdam, Netherlands, for providing us *E.faecium* W54 and *L.brevis* W63, as well as their sequencing data; Dr. Jan Kok of Department of Molecular genetics, University of Groningen, Netherlands, and Dr. Miguel A. Alvarez of Instituto de Productos Lácteos de Asturias, Villaviciosa, Spain for providing the mutant strain *E. faecalis* v583; Dr. Uwe Tietge and Rema H. Mistry, MSc. of the Department of Pediatrics, University Medical Centre Groningen, Groningen, Netherlands for providing assistance with germ-free and specific pathogen free mice; and Dr. Phillip A. Engen, Division of Digestive Disease and Nutrition, Section of Gastroenterology, Rush University Medical Center, USA for assisting in preparing fecal samples from Parkinson’s patients for shipment.

## Authors Contributions

S.P.K. and S.E.A conceived and designed the study. S.P.K, A.K.F., A.O.E.G., M.C., A.K., G.D., S.E.A performed the experiments and S.P.K and S.E.A analyzed the data. S.P.K and S.E.A. wrote the original manuscript that was reviewed by A.K.F., S.E.A., A.K., G.D. Funding was acquired by S.E.A.

## STAR-methods

### EXPERIMENTAL MODEL AND SUBJECT DETAILS

#### Human fecal samples from patients with Parkinson’s disease

Fecal samples from patients diagnosed with Parkinson’s disease (n=10) on variable doses (300-1100mg L-DOPA per day) of L-DOPA/carbidopa treatment were acquired from the Movement Disorder Center at Rush University Medical Center, Chicago, Illinois, USA. Patients’ characteristics were published previously (Keshavarzian et al., 2015) (more details are provided in Table S4). Solid fecal samples were collected in anaerobic fecal bags and kept sealed in a cold environment until brought to the hospital where they were immediately stored at −80°C until analysis.

#### Rats

Twenty-five male wild-type Groningen rats ((Koolhaas et al., 2013); Groningen breed, male, age 18-24 weeks) housed 4-5 animals/cage had *ad libitum* access to water and food (RMH-B, AB Diets; Woerden, the Netherlands) in a temperature (21 ± 1°C) and humidity-controlled room (45–60% relative humidity), with a 12 hrs light/dark cycle (lights off at 1:00 p.m.). On ten occasions over a period of three weeks, rats were taken from their social housing cage between circadian times 6 and 16.5, and put in an individual training cage (L×W×H = 25×25×40 cm) with a layer of their own sawdust without food and water. Ten minutes after transfer to these cages, rats were offered a drinking pipet in their cages with a 2.5 ml saccharine-solution (1.5 g/L, 12476, Sigma). Over the course of training, all rats learned to drink the saccharine-solution avidly. On the 11^th^ occasion, the saccharine solution was used as vehicle for the L-DOPA/carbidopa mixture (15/3.75 mg/kg), which all rats drank within 15 seconds. Fifteen minutes after drinking the latter mixture (maximum bioavailability time point of L-DOPA in blood as previously described (*18*), the rats were anesthetized with isoflurane and sacrificed. Blood was withdrawn by heart puncture and placed in tubes pre-coated with 5 mM EDTA. The collected blood samples were centrifuged at 1500 x g for 10 minutes at 4°C and the plasma was stored at −80°C prior to L-DOPA, dopamine, and DOPAC extraction. Luminal contents were harvested from the rat jejunum by gentle pressing and were snap frozen in liquid N_2_, stored at −80°C until used for qPCR, and extraction of L-DOPA and its metabolites. Oral administration (by drinking, with saccharine as vehicle) of L-DOPA was corrected for by using carbidopa as an internal standard to correct for intake. Further, 5 rats were used as control and were administered a saccharine only solution (vehicle) to check for basal levels of L-DOPA, dopamine, and DOPAC levels or background HPLC-peaks. Jejunal content of control rats was used in *ex vivo* fermentation experiments (see incubation experiments of jejunal content section). All animal procedures were approved by the Groningen University Committee of Animal experiments (approval number: AVD1050020184844), and were performed in adherence to the NIH Guide for the Care and Use of Laboratory Animals.

#### Bacteria

*Escherichia coli* DH5a or BL21 were routinely grown aerobically in Luria-Broth (LB) at 37°C degrees with continuous agitation. Other strains listed in **Table S5** were grown anaerobically (10% H_2_, 10% CO_2_, 80% N_2_) in a Don Whitley Scientific DG250 Workstation (LA Biosystems, Waalwijk, The Netherlands) at 37°C in an enriched beef broth based on SHIME medium (Auchtung et al., 2015) (**Table S6**). Bacteria were inoculated from −80°C stocks and grown overnight. Before the experiment, cultures were diluted 1:100 in fresh medium from overnight cultures. L-DOPA (D9628, Sigma, The Netherlands), carbidopa (C1335, Sigma), benserazide (B7283, Sigma), or methyldopa (857416, Sigma) were supplemented during the lag or stationary phase depending on the experiment. Growth was followed by measuring the optical density (OD) at 600 nM in a spectrophotometer (UV1600PC, VWR International, Leuven, Belgium).

### METHOD DETAILS

#### Cloning and heterologous gene expression

The human DOPA decarboxylase gene cloned in pET15b was ordered from GenScript (Piscataway, USA) (**Table S5**). Tyrosine decarboxylase-encoding genes from *E. faecalis* v583 (TDC_EFS,_ accession: EOT87933), *E. faecium* W54 (TDC_EFM_, accession: MH358385; ^P^TDC_EFM_, accession: MH358384) were amplified using Phusion High-fidelity DNA polymerase and primers listed in **Table S7**. All amplified genes were cloned in pET15b, resulting in pSK18, pSK11, and pSK22, respectively (**Table S5**). Plasmids were maintained in *E. coli* DH5α and verified by Sanger sequencing before transformation to *E. coli* BL21 (DE3). Overnight cultures were diluted 1:50 in fresh LB medium with the appropriate antibiotic and grown to OD_600_ = 0.7-0.8. Protein translation was induced with 1mM Isopropyl β-D-1-thiogalactopyranoside (IPTG, 11411446001, Roche Diagnostics) and cultures were incubated overnight at 18°C. Cells were washed with 1/5th of 1× ice-cold PBS and stored at −80 °C or directly used for protein isolation. Cell pellets were thawed on ice and resuspended in 1/50th of buffer A (300 mM NaCl; 10 mM imidazole; 50 mM KPO4, pH 7.5) containing 0.2 mg/mL lysozyme (105281, Merck) and 2 µg/mL DNAse (11284932001, Roche Diagnostics), and incubated for at least 10 minutes on ice before sonication (10 cycles of 15s with 30s cooling at 8 microns amplitude) using Soniprep-150 plus (Beun de Ronde, Abcoude, The Netherlands). Cell debris were removed by centrifugation at 20000 × *g* for 20 min at 4°C. The 6×his-tagged proteins were purified using a nickel-nitrilotriacetic acid (Ni-NTA) agarose matrix (30250, Qiagen). Cell-free extracts were loaded on 0.5 ml Ni-NTA matrixes and incubated on a roller shaker for 2 hours at 4°C. The Ni-NTA matrix was washed three times with 1.5 ml buffer B (300 mM NaCl; 20 mM imidazole; 50 mM KPO4, pH 7.5) before elution with buffer C (300 mM NaCl; 250 mM imidazole; 50 mM KPO4, pH 7.5). Imidazole was removed from purified protein fractions using Amicon Ultra centrifugal filters (UFC505024, Merck) and washed three times and reconstituted in buffer D (50 mM Tris-HCL; 300 mM NaCl; pH 7.5) for TDC_EFS_, and TDC_EFM,_ buffer E (100 mM KPO4; pH 7.4) for^P^TDC_EFM_ and buffer F (100 mM KPO4; 0.1 mM pyridoxal-5-phosphate; pH 7.4) for DDC. Protein concentrations were measured spectrophotometrically (Nanodrop 2000, Isogen, De Meern, The Netherlands) using the predicted extinction coefficient and molecular weight from ExPASy ProtParam tool (www.web.expasy.org/protparam/).

#### Enzyme kinetics and IC50 curves

Enzyme kinetics were performed in 200 mM potassium acetate buffer containing 0.1 mM PLP (pyridoxal-5-phosphate, P9255, Sigma, The Netherlands) and 10 nM of enzyme at pH 5 for TDC_EFS_ and TDC_EFM_, and pH 4.5 for ^P^TDC_EFM_. Reactions were performed in triplicate using L-DOPA substrate ranges from 0.5-12.5 mM and tyrosine substrate ranges from 0.25- 2.5 mM. Michaelis-Menten kinetic curves were fitted using GraphPad Prism 7. The human dopa decarboxylase kinetic reactions were performed in 100 mM potassium phosphate buffer at pH 7.4 containing 0.1 mM PLP and 10 nM enzyme concentrations with L-DOPA substrate ranges from 0.1-1.0 mM. Reactions were stopped with 0.7% HClO_4_, filtered and analyzed on the HPLC-ED-system described below. For IC50 curves, the reaction was performed using L-DOPA as the substrate at concentrations lower or equal to the Km of the decarboxylases (DDC, 0.1 mM; TDC_EFS_ and TDC_EFM_, 1.0 mM; ^P^TDC_EFM_, 0.5 mM) with 10 different concentrations of carbidopa in triplicate (human dopa decarboxylase, 0.005-2.56 µM; bacterial tyrosine decarboxylases, 2-1024 µM).

#### HPLC-ED analysis and sample preparation

1 mL of ice-cold methanol was added to 0.25 mL cell suspensions. Cells and protein precipitates were removed by centrifugation at 20000 × *g* for 10 min at 4°C. Supernatant was transferred to a new tube and the methanol fraction was evaporated in a Savant speed-vacuum dryer (SPD131, Fisher Scientific, Landsmeer, The Netherlands) at 60°C for 1h 15 min. The aqueous fraction was reconstituted to 1 mL with 0.7% HClO_4_. Samples were filtered and injected into the HPLC system (Jasco AS2059 plus autosampler, Jasco Benelux, Utrecht, The Netherlands; Knauer K-1001 pump, Separations, H. I. Ambacht, The Netherlands; Dionex ED40 electrochemical detector, Dionex, Sunnyvale, USA, with a glassy carbon working electrode (DC amperometry at 1.0 V or 0.8 V, with Ag/AgCl as reference electrode)). Samples were analyzed on a C18 column (Kinetex 5µM, C18 100 Å, 250 ×4.6 mm, Phenomenex, Utrecht, The Netherlands) using a gradient of water/methanol with 0.1% formic acid (0–10 min, 95–80% H_2_O; 10-20 min, 80-5% H_2_O; 20-23 min 5% H_2_O; 23-31 min 95% H_2_O). Data recording and analysis was performed using Chromeleon software (version 6.8 SR13).

#### Bioinformatics

TDC_EFS_ (NCBI accession: EOT87933) was BLASTed against the protein sequences from the NIH HMP data bank using search limits for Entrez Query “43021[BioProject]”. All BLASTp hits were converted to a distance tree using NCBI TreeView (Parameters: Fast Minimum Evolution; Max Seq Difference, 0.9; Distance, Grishin). The tree was exported in Newick format and visualized in iTOL phylogentic display tool (http://itol.embl.de/). Whole genomes, or contigs containing the TDC gene cluster were extracted from NCBI and aligned using Mauve multiple genome alignment tool (v 2.4.0, www.darlinglab.org/mauve/mauve.html).

#### Incubation experiments of jejunal content

Luminal contents from the jejunum of wild-type Groningen rats (n=5) were suspended in EBB (5% w/v) containing 1 mM L-DOPA and incubated for 24 hours in an anaerobic chamber at 37 °C prior to HPLC-ED analysis (DC amperometry at 0.8 V).

#### DNA extraction

DNA was extracted from fecal samples of Parkinson’s patients and jejunal contents of rats using QIAGEN (Cat no. 51504) kit-based DNA isolation as previously described (Zoetendal et al., 2006) with the following modifications: fecal samples were suspended in 1 mL inhibitEX buffer (1:5 w/v) and transferred to screw-caped tubes containing 0.5 g of 0.1 mm and 3 mm glass beads. Samples were homogenized 3 × 30 sec with 1-minute intervals on ice in a mini bead-beater (Biospec, Bartlesville, USA) 3 times.

#### Quantification of bacterial tyrosine decarboxylase

To identify bacterial species carrying the *tdc* gene, a broad range of *tdc* genes from various bacterial genera were targeted as previously described (Torriani et al., 2008) (**Figure S5**). Quantitative PCR (qPCR) of *tdc* genes was performed on DNA extracted from each fecal sample of Parkinson’s patients and rats’ jejunal content using primers (Dec5f and Dec3r) targeting a 350bp region of the *tdc* gene. Primers targeting 16sRNA gene for all bacteria (Eub338 and Eub518) were used as an internal control (**Table S7**). All qPCR experiments were performed in a Bio-Rad CFX96 RT-PCR system (Bio-Rad Laboratories, Veenendaal, The Netherlands) with iQ SYBR Green Supermix (170-8882, Bio-Rad) in triplicate on 20 ng DNA in 10 µL reactions using the manufacturer’s protocol. qPCR was performed using the following parameters: 3 min at 95°C; 15 sec at 95°C, 1 min at 58°C, 40 cycles. A melting curve was determined at the end of each run to verify the specificity of the PCR amplicons. Data analysis was performed using the BioRad software. Ct[DEC] values were corrected with the internal control (Ct[16s]) and linearized using 2^-(Ct[DEC]-Ct[16s]) based on the 2^-ΔΔCt method (Livak and Schmittgen, 2001).

#### Jejunal and plasma extraction of L-DOPA and its metabolites

L-DOPA, dopamine, and DOPAC were extracted from each luminal jejunal content and plasma samples of rats using activated alumina powder (199966, Sigma) as previously described (Ganhao et al., 1991) with a few modifications. 50-200 µl blood plasma was used with 1µM DHBA (3, 4-dihydroxybenzylamine hydrobromide, 858781, Sigma) as an internal standard. For jejunal luminal content samples, an equal amount of water was added (w/v), and suspensions were vigorously mixed using a vortex. Suspensions were subsequently centrifuged at 20000 × *g* for 10 min at 4°C. 50-200 µL of supernatant was used for extraction. Samples were adjusted to pH 8.6 with 200–800µl TE buffer (2.5% EDTA; 1.5 M Tris/HCl pH 8.6) and 5-10 mg of alumina was added. Suspensions were mixed on a roller shaker at room temperature for 15 min and were thereafter centrifuged for 30s at 20000 × *g* and washed twice with 1 mL of H_2_O by aspiration. L-DOPA and its metabolites were eluted using 0.7% HClO_4_ and filtered before injection into the HPLC-ED-system as described above (DC amperometry at 0.8 V).

### QUANTIFICATION AND STATISTICAL ANALYSIS

All statistical tests and (non)linear regression models were performed using GraphPad Prism 7. Statistical tests performed are unpaired T-tests, 2-way-ANOVA followed by a Fisher’s LSD test. Specific tests and significance are indicated in the figure legends.

## Supplementary material

### Supplementary methods

#### Screening for *Enterococcus* strains, isolated from healthy subjects and clinical isolates

*Enterococcus* strains were isolated from fecal or urine samples from clinical setting or from healthy volunteers (aged 2 to 79 years). All samples were collected between January 2008 and 2018 in Beni-Suef City, Egypt. A total of 77 *Enterococcus* spp. were isolated on bile esculin agar and observed microscopically after gram staining. The screening for decarboxylase activity was performed as described previously by Bover-Cid and Holzapfel (Bover-Cid and Holzapfel, 1999) with L-DOPA or tyrosine added as a substrate to a final concentration of 1% to screen for production of dopamine and tyramine. All *Enterococcus* strains were spot inoculated on agar plates containing the substrates of interest and on control plates without any substrate. The plates were duplicated, either aerobically or anaerobically at 37°C. Plates were checked daily for 4 days to record the change in the indicator color from yellow to violet, indicative of production of tyramine and dopamine, respectively (**Table S1**).

#### Fecal samples from patients with Parkinson’s disease

All study subjects consented to the use of their samples for research. Parkinson’s disease was diagnosed according to the UK Brain Bank Criteria. Exclusion criteria for Parkinson’s disease subjects: (1) atypical or secondary Parkinsonism, (2) the use of probiotics or antibiotics within three months prior to sample collection, (3) primary gastrointestinal pathology, (4) unstable medical, neurological, or psychiatric illness, (5) low platelet count (<80k), uncorrectable prolonged PT (>15 sec. Solid fecal samples were collected via a home feces collection kit. Study patients were provided with the supplies and instructions for home feces collection using the BD Gaspak EZ Anaerobe Gas Generating Pouch System with Indicator (Ref 260683; Becton, Dickinson and Company, Sparks, MD) in order to minimize the exposure of the feces to high oxygen ambient atmosphere, which may alter the microbiota. Patients were asked to have a bowel movement within 24 hours of their study visit. Patients kept the sealed anaerobic fecal bag in a cold environment, before bringing the anaerobic fecal bag to the hospital. Fecal samples were then immediately stored at −80°C until analysis.

#### Oral administration of L-DOPA to germ-free and specific pathogen free mice

Specific pathogen free (SPF) C57BL/6OlaHsd 8-week old females were purchased from a commercial supplier (Envigo, Horst, the Netherlands), and C57BL/6OlaHsd 8-week old germ-free (GF) female mice were obtained from a breeding colony at the animal facility at the Radboud University (Nijmegen, the Netherlands). Animals were receiving a D12450B diet (10% fat, Research Diet Services, Wijk bij Duurstede, the Netherlands). Mice (n = 3/group) were acclimatized for two weeks and were administered 15 mg/kg L-DOPA freshly dissolved in tap water by oral gavage. Before and 15 min after oral-gavage, blood samples were collected retro-orbitally with heparin-coated capillaries. The collected blood samples were centrifuged at 2000 x g for 15 minutes at 4°C and the plasma was stored at −80°C prior to catechol extraction. Experiments were approved by the animal ethics committee of the University of Groningen (Dieren experimenten commissie Rijksuniversiteit Groningen, DEC-RUG).

#### Primers targeting bacterial tyrosine decarboxylase

In order to cover numerous bacterial species harboring the *tdc* gene, degenerate primers, Dec5f and Dec3r, previously designed to target 350bp region of the *tdc* gene from a variety of bacterial genera (Torriani et al., 2008) were used for qPCR. Prior to qPCR experiments a normal PCR test on the genomic DNA *E. faecalis* v583 was performed using Phire Hot Start II DNA Polymerase (F125S, ThermoFisher) using the following PCR parameters: 3 min at 98°C; 15 sec at 98°C, 1 min at 58°C and 1 min at 72°C, for 35 cycles; 5 min at 72°C, to ensure primer specificity.

## Supplementary Figures, Tables, and Legends

**Figure S1.**
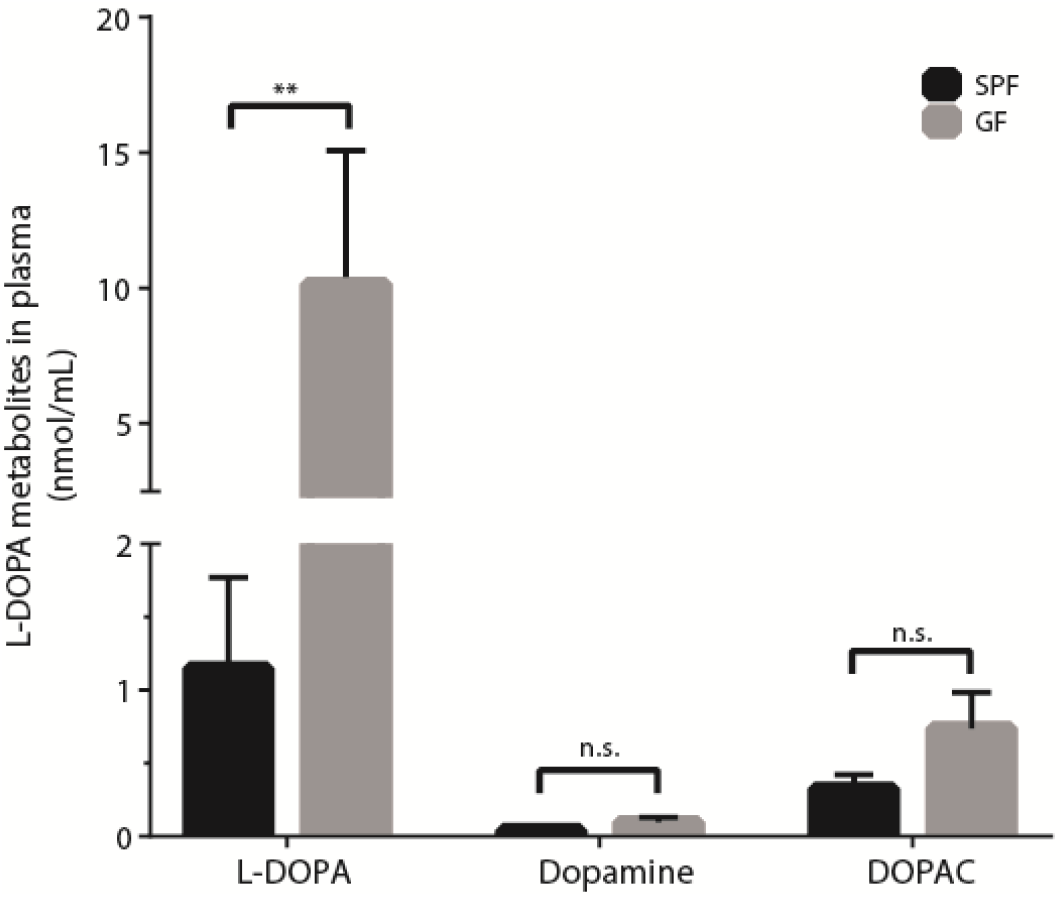
Related to Figure 1. L-DOPA plasma levels are higher in germ-free mice compared to specific pathogen-free mice. Bar graphs showing the mean levels of L-DOPA, dopamine, and DOPAC (3,4-Dihydroxyphenylacetic acid) in plasma of germ-free (GF, n=3) and specific pathogen free mice (SPF, n=3). Levels of L-DOPA and its metabolite DOPAC are significantly higher in GF-mice. Error bars represent SEM and significance was tested using a 2-way ANOVA followed by Fisher’s LSD test (**=*p<0.01*).

**Figure S2.**
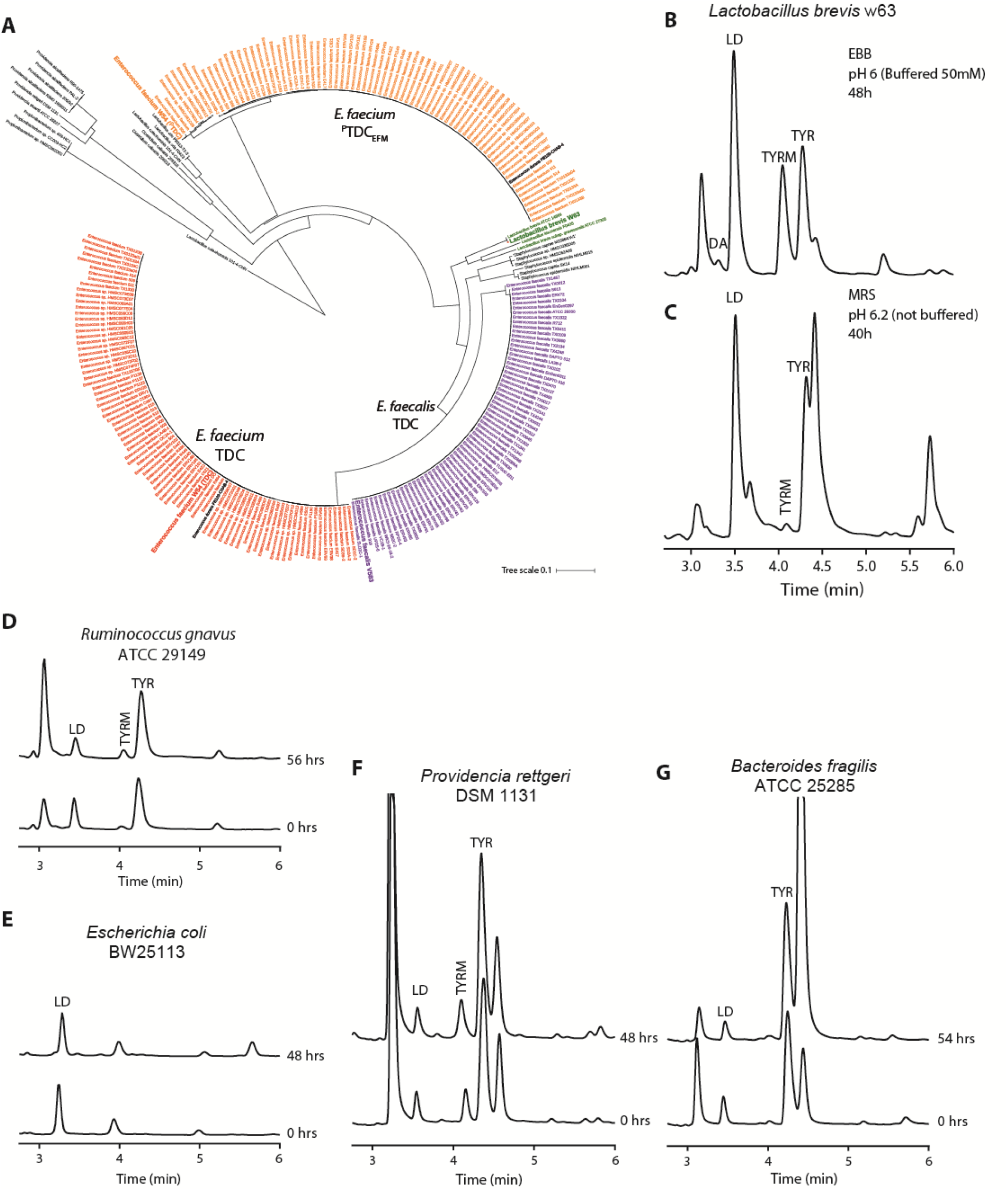
Related to Figure 2. Microbiota harboring other PLP-dependent amino acid decarboxylases do not decarboxylate L-DOPA. (**A**) Phylogenetic analysis of TDC proteins from NIH Human Microbiome Project (HMP) protein database, TDC_EFS_ (EOT87933) was used as query. TDC protein sequences from strains employed in this study are depicted in bold. Live stationary cultures of *L. brevis* grown with L-DOPA in (**B**) MRS (De Man, Rogosa and Sharpe) or in (**C**) enriched beef broth (EBB) buffered at pH 6.0. (**D-G**) Gut associated bacteria harboring different amino acid decarboxylases, which were previously identified (*38*) (**Table S1**) were tested for their ability to convert L-DOPA in live stationary cultures.

**Figure S3.**
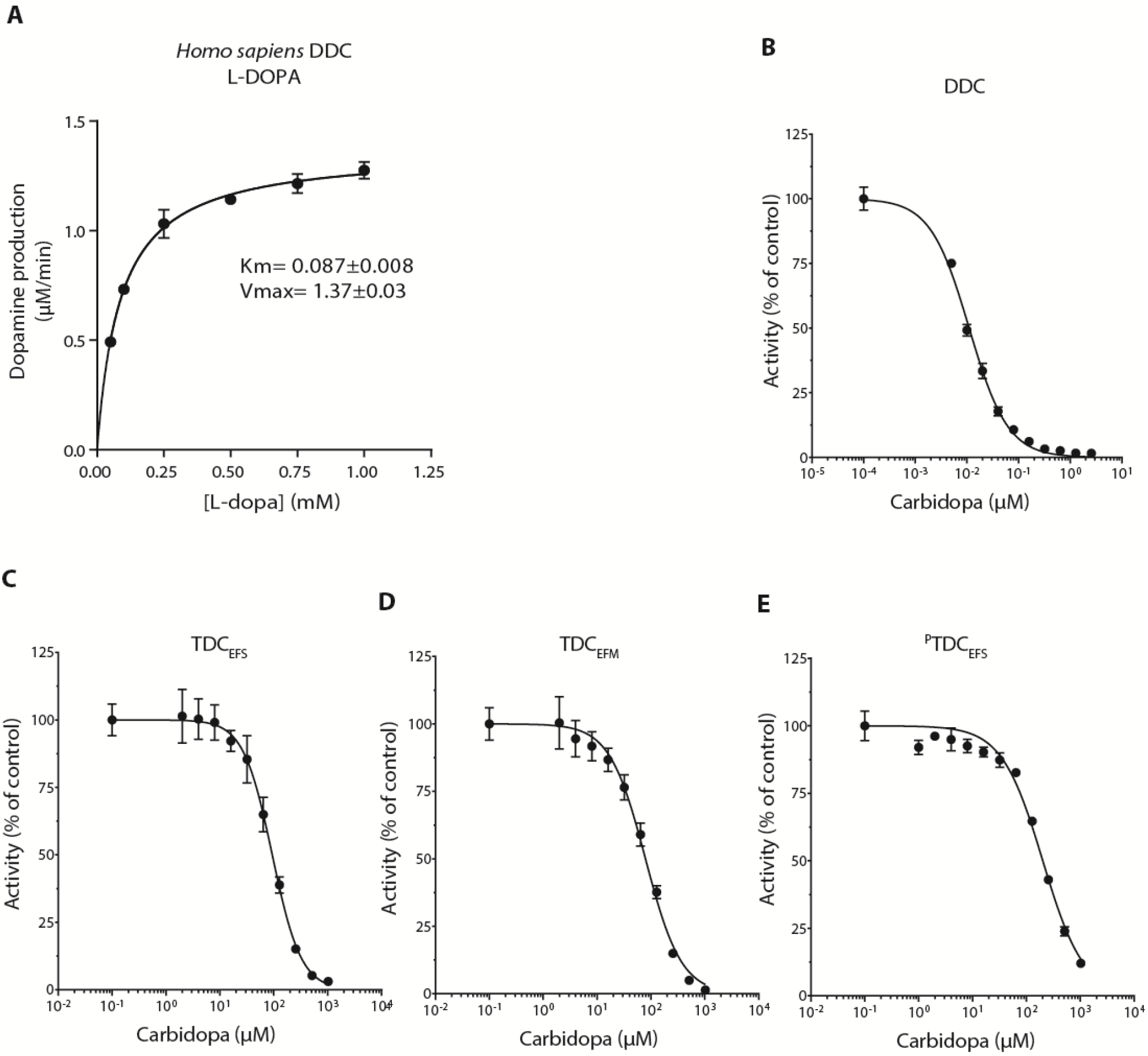
Related to Figure 4. IC50 determination for human DOPA decarboxylase and bacterial tyrosine decarboxylases. (**A**) Kinetic curve with L-DOPA as substrate for human DOPA decarboxylase (DDC) to determine the inhibitory constant for carbidopa. Reactions were performed in triplicate using 10 nM of enzyme in 100 mM PO_4_ with 100 µM PLP (Montioli et al., 2011) with L-DOPA concentrations ranging from 0.1-1.0 mM. The enzyme kinetic parameters were calculated using nonlinear Michaelis-Menten regression model (for further kinetic parameters see **Table 1**). IC50 inhibitory curves using carbidopa as inhibitor for (**B**) DDC (0.005-2.56 µM carbidopa), (**C**) TDC_EFS_ (2-1024 µM carbidopa), (**D**) TDC_EFM_ (2-1024 µM carbidopa), (**E**) ^P^TDC_EFM_ (2-1024 µM carbidiopa). Reactions were performed in triplicate and the parameters were determined by fitting a sigmoidal–curve ([inhibitor] vs. normalized response). Further parameters are listed in **Table S3**.

**Figure S4.**
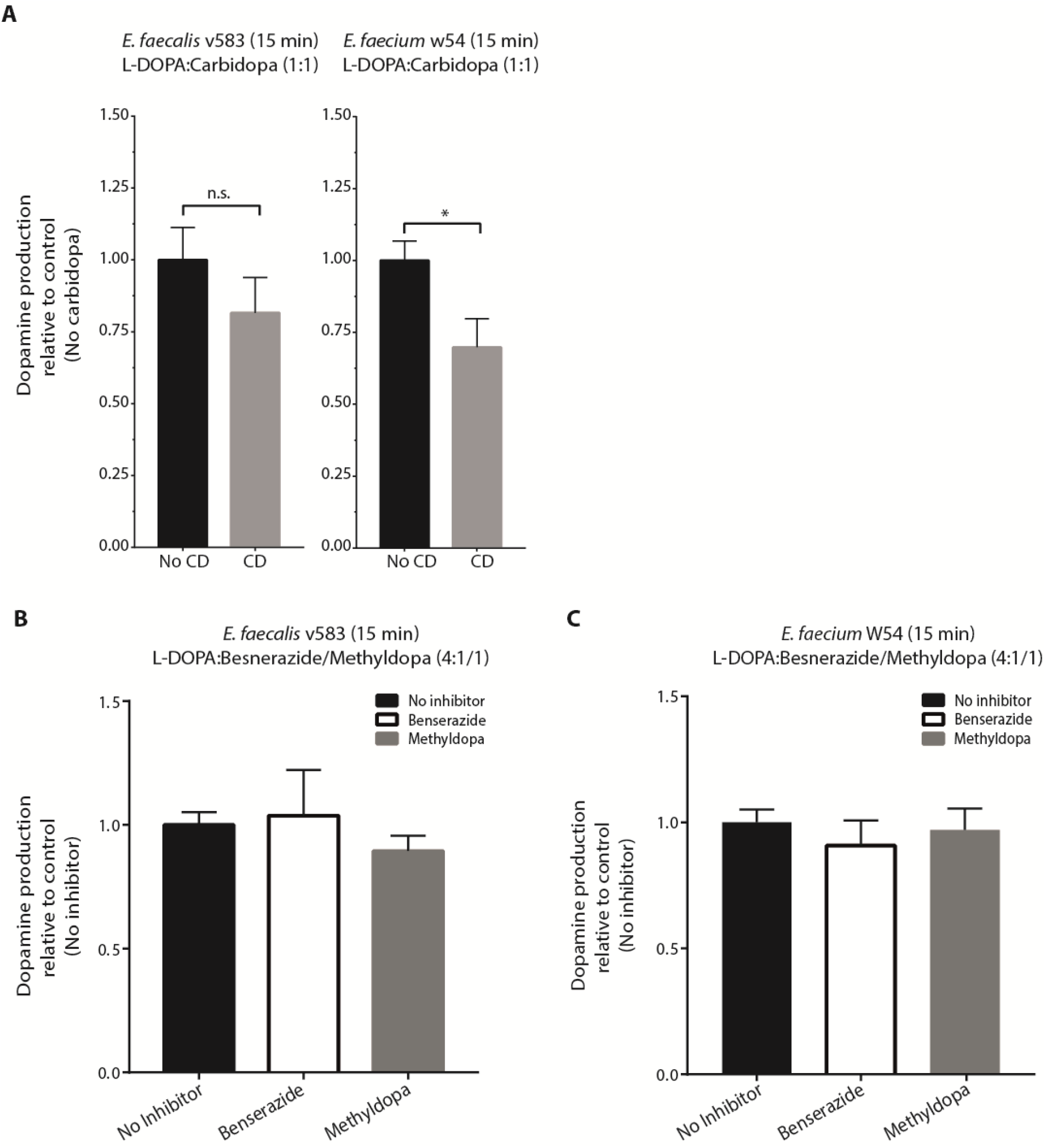
Related to Figure 4. Human DOPA decarboxylase inhibitors are ineffective against the decarboxylase activity of live enterococci. Live stationary cultures of *E. faecalis* and *E. faecium* incubated for 15 minutes with (**A**) an equimolar ratio of L-DOPA/carbidopa, (**B, C**) a 6:1 molar ratio (4:1 in weight) of L-DOPA/benserazide or 4.8:1 molar ratio (4:1 in weight) of L-DOPA/methyldopa (100/16.7/20,8 µM L-DOPA/benserazide/methyldopa). Samples were analyzed using HPLC-ED. Bar graphs show levels of dopamine production (relative to control, where no inhibitor was added) with and without the addition of inhibitor. Error bars represent the SEM and significance was tested using a parametric unpaired T-test (*=p<0.02).

**Figure S5.**
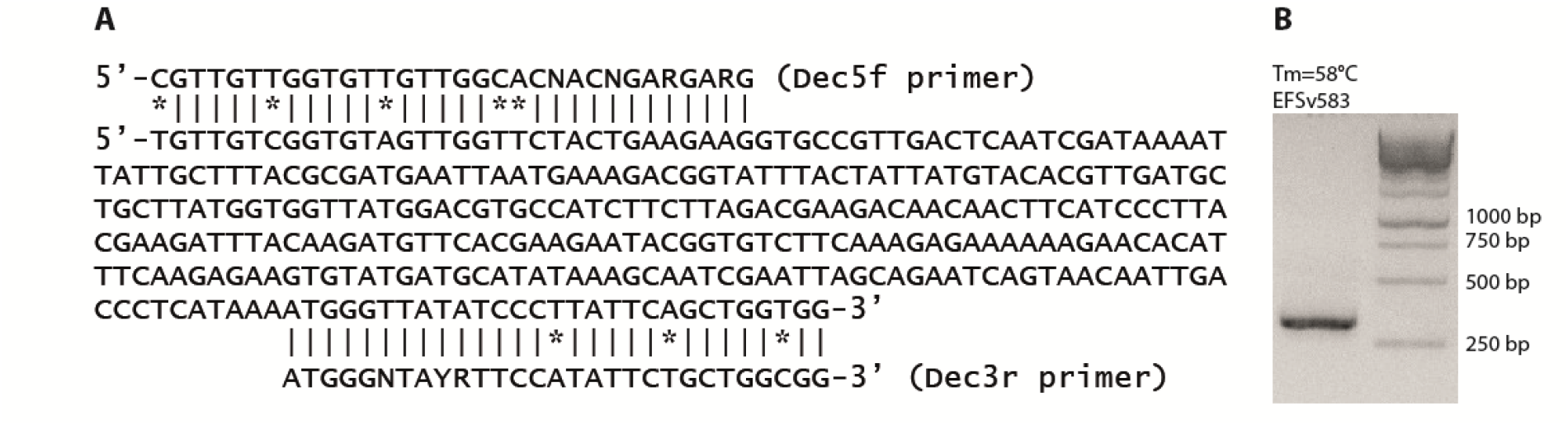
Related to Figure 5 and 6. Primers (Dec5f and Dec3r) targeting *E. faecalis* v583 *tdc* gene. (**A**) Target of the Dec5f and Dec3r primers (Torriani et al., 2008) are depicted for *Enterococcus faecalis* v583 with (**B**) the corresponding agarose gel of PCR amplification of 336 bp fragment of *tdc-*gene.

**Table S1.** Related to Figure 2. Enterococcus strains isolated from healthy subjects and clinical enterococcal isolates. *Enterococcus* strains isolated from (1) fecal samples (collected from non-hospitalized subjects from clinical labs during their routine check up), (2) urine samples (collected from non-hospitalized patients suffering from urinary tract infection), and (3) healthy volunteers (age 2 to 79 years). All samples were isolated between January 2008- 2018 in Beni-Suef City, Egypt. 72 out of 77 isolates were able to decarboxylate L-DOPA and tyrosine indicating that only 5 isolates are species or strains not encoding for tyrosine decarboxylase.

|  | Number | Sample | Type | Bile<br>aesculin<br>hydrolysis | L-DOPA<br>decarboxylation | Tyrosine<br>decarboxylation |
| --- | --- | --- | --- | --- | --- | --- |
|  | 29 | Clinical | Stool | 29 | 29 | 29 |
|  | 34 | Clinical | Urine | 34 | 33 | 33 |
|  | 14 | Volunteer | Stool | 14 | 10 | 10 |
| **Total** | **77** |  |  | **77** | **72** | **72** |

**Table S2.** Related to Figure 2. List of microbiota harboring other PLP-dependent amino acid decarboxylases tested in this study. * Protein identity from strain tested compared to cloned gene Williams et al. (Williams et al., 2014). NA, not applicable.

| Reported tyrosine decarboxylation (3) | Cloned gene (3) | Associated protein accession | Definition | This study (Live bacterial culture) | Protein Identity* | Protein accession |
| --- | --- | --- | --- | --- | --- | --- |
| - | BF 0393 (BF9343_0382) | CAH06161 | Putative Glutamate decarboxylase | Bacteroides fragilis NCTC 9343 (=ATCC 25285) | 100.00% | CAH06161 |
|  | Not tested | NA | Tyrosine decarboxylase | Enterococcus faecalis v583 | NA | EOT87933 |
|  | Not tested | NA | Tyrosine decarboxylase | Enterococcus faecium W54 | NA | MH358384, MH358385 |
| - | ECD_03365 | ACT45166 | Glutamate decarboxylase | Escherichia coli BW25113 | 100.00% | AIN33843 |
| - | LVIS_1847 | ABJ64910 | Glutamate decarboxylase | Lactobacillus brevis W63 | NA |  |
| ++ | LVIS_2213 | ABJ65263 | Tyrosine decarboxylase | Lactobacillus brevis W63 | 100.00% | MH3583846 |
| - | LVIS_0079 | ABJ63253 | Glutamate decarboxylase | Lactobacillus brevis W63 | NA |  |
| +++ | PROSTU_02509 | EDU59320 | putative tyrosine decarboxylase | Providencia rettgeri DSM1131 | 83% | EFE55437 |
| +++ | RUMGNA_01526 | EDN78222 | Tryptophan decarboxylase | Ruminococcus gnavus ATCC29149 | 100.00% | EDN78222 |

**Table S3.** Related to Figure 4. IC50 curve parameters. The parameters were determined by fitting a sigmoidal–curve ([inhibitor] vs. normalized response) using Graphpad Prism. Reactions were performed in triplicate.

|  | **DDC** | **TDC<sub>EFS</sub>** | **TDC<sub>EFM</sub>** | **PTDC<sub>EFM</sub>** |
| --- | --- | --- | --- | --- |
| **[E] (nM)** | 10 | 10 | 10 | 10 |
| **[S] (μM)** | 100 | 1000 | 1000 | 500 |
| **[Km] (μM)** | 87.3 | 3043.0 | 7244.0 | 448.1 |
| **HillSlope** | 1.16 | 1.6 | 1.3 | 1.1 |
| **IC50 (μM)** | 0.01 | 93.8 | 79.2 | 201.3 |
| **Ki=IC50/(1+([S]/Km))** | 0.005 | 70.6 | 69.6 | 95.1 |
| **[IC50]/[s]** | 0.01% | 9% | 8% | 40% |

**Table S4.**
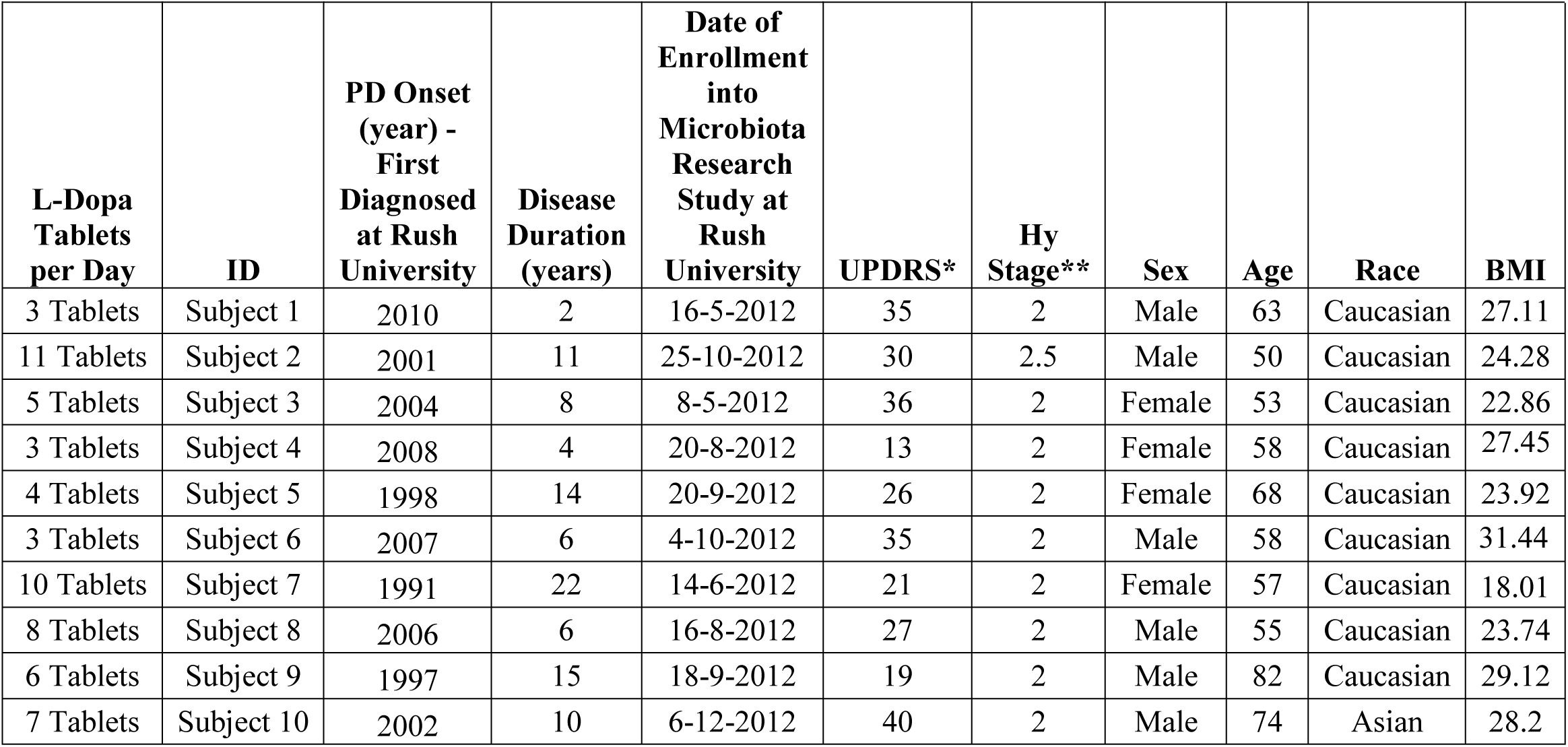
Related to Figure 5. Sample information, Parkinson’s patients. * Unified Parkinson Disease Rating Scale, ** Hoehn and Yahr Stage, BMI = Body Mass Index. According to medical records, all subjects were on the same L-DOPA dose thereafter their PD Onset (year) at Rush University Medical Center Neurology Department. Parkinson’s disease was diagnosed according to the UK Brain Bank Criteria (Keshavarzian et al., 2015).

**Table S5.**
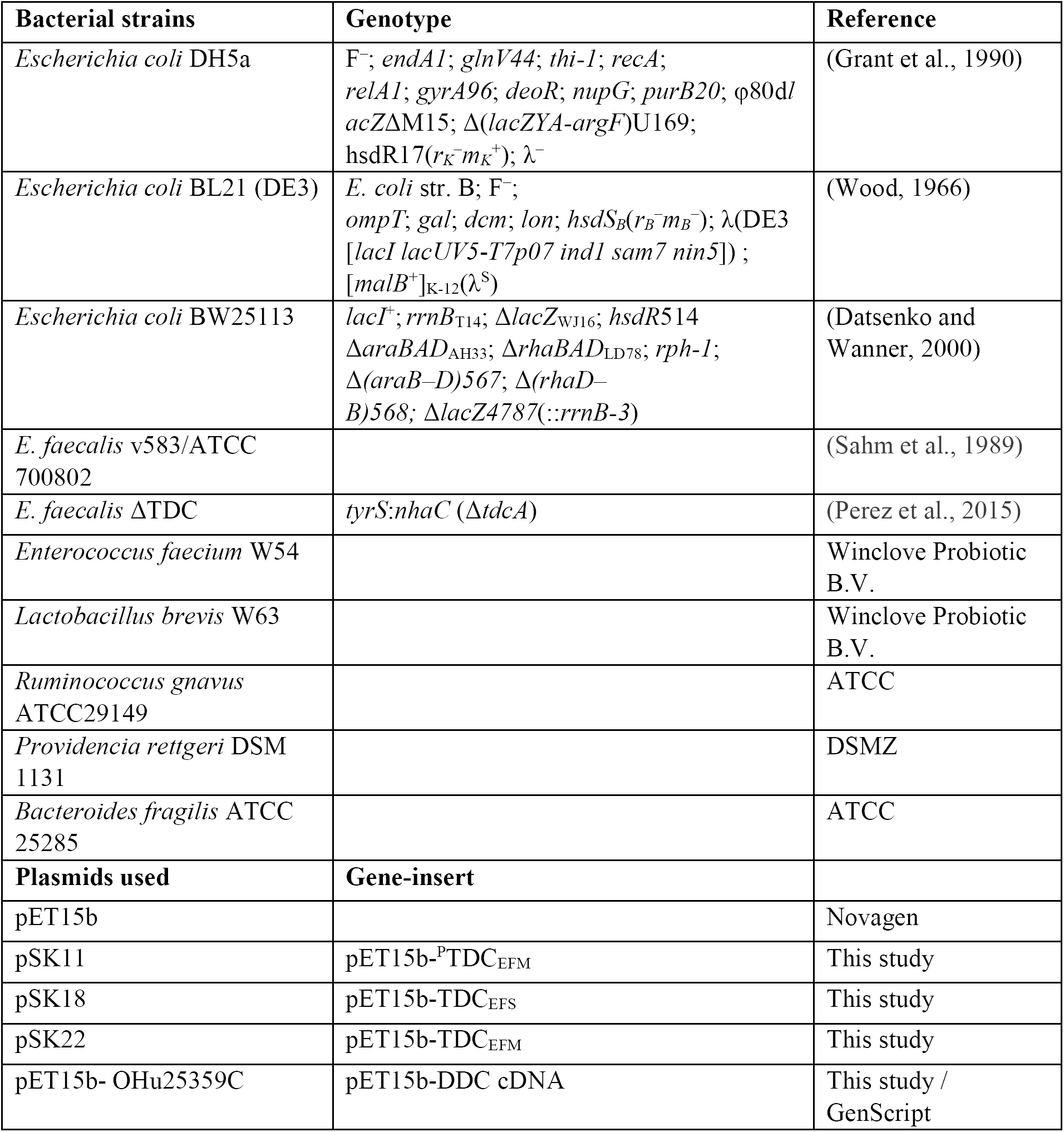
Bacterial strains and Plasmids used in this study.

| Bacterial strains | Genotype | Reference |
| --- | --- | --- |
| *Escherichia coli* DH5a | F<sup>-</sup>; *endA1*; *glnV44*; *thi-1*; *recA*; *relA1*; *gyrA96*; *deoR*; *nupG*; *purB20*; $\phi 80dl$ *acZ*ΔM15; $\Delta$(*lacZYA-argF*)U169; *hsdR17*(*r<sub>K</sub><sup>-</sup>m<sub>K</sub><sup>+</sup>*); $\lambda^-$ | (Grant et al., 1990) |
| *Escherichia coli* BL21 (DE3) | *E. coli* str. B; F<sup>-</sup>; *ompT*; *gal*; *dcm*; *lon*; *hsdS<sub>B</sub>*(*r<sub>B</sub><sup>-</sup>m<sub>B</sub><sup>-</sup>*); $\lambda$(DE3 [*lacI lacUV5-T7p07 ind1 sam7 nin5*]) ; [*malB<sup>+</sup>*]<sub>K-12</sub>($\lambda^S$) | (Wood, 1966) |
| *Escherichia coli* BW25113 | *lacI<sup>+</sup>*; *rrnB<sub>T14</sub>*; Δ*lacZ<sub>WJ16</sub>*; *hsdR514* Δ*araBAD<sub>AH33</sub>*; Δ*rhaBAD<sub>LD78</sub>*; *rph-1*; $\Delta$(*araB-D*)567; $\Delta$(*rhaD-B*)568; Δ*lacZ4787*(::*rrnB-3*) | (Datsenko and Wanner, 2000) |
| *E. faecalis* v583/ATCC 700802 |  | (Sahm et al., 1989) |
| *E. faecalis* ΔTDC | *tyrS:nhaC* (Δ*tdcA*) | (Perez et al., 2015) |
| *Enterococcus faecium* W54 |  | Winclove Probiotic B.V. |
| *Lactobacillus brevis* W63 |  | Winclove Probiotic B.V. |
| *Ruminococcus gnavus* ATCC29149 |  | ATCC |
| *Providencia rettgeri* DSM 1131 |  | DSMZ |
| *Bacteroides fragilis* ATCC 25285 |  | ATCC |
| Plasmids used | Gene-insert |  |
| pET15b |  | Novagen |
| pSK11 | pET15b-<sup>P</sup>TDC<sub>EFM</sub> | This study |
| pSK18 | pET15b-TDC<sub>EFs</sub> | This study |
| pSK22 | pET15b-TDC<sub>EFM</sub> | This study |
| pET15b- OHu25359C | pET15b-DDC cDNA | This study / GenScript |

**Table S6.** Constituents of enriched beef broth (EBB) medium used in this study. The KPO_4_ solution is buffered at 50 mM, pH7.

| **Component** | **g/L** |
| --- | --- |
| Glucose | 2.000 |
| NaCl | 0.080 |
| $\text{K}_2\text{HPO}_4$ | 5.310 |
| $\text{KH}_2\text{PO}_4$ | 2.650 |
| $\text{NaHCO}_3$ | 0.400 |
| Beef extract | 5.000 |
| Yeast extract | 3.000 |
| Peptone | 0.600 |
| $\text{CaCl}_2$ | 0.008 |
| $\text{MgSO}_4$ | 0.008 |
| Cysteine | 0.500 |
| Hemin | 0.005 |
| **Vitamine solution (1000x)** |  |
| D-biotin | 0.0020 |
| D-Pantothenic acid | 0.0100 |
| Ca2.Nicotinamide | 0.0050 |
| Vitamin B12 | 0.0005 |
| Thiamin.HCl | 0.0040 |
| Para-aminobenzoic acid | 0.0050 |
| Riboflavin | 0.0050 |
| Folic acid | 0.0020 |
| Pyridoxyl-5-Phosphate | 0.0100 |
| Vitamin K1 | 0.0005 |
| **Trace Elements (1000x)** |  |
| EDTA | 1.000 |
| $\text{ZnSO}_4 \cdot 7\text{H}_2\text{O}$ | 0.178 |
| $\text{MnSO}_4 \cdot \text{H}_2\text{O}$ | 0.452 |
| $\text{FeSO}_4 \cdot 7\text{H}_2\text{O}$ | 0.100 |
| $\text{CoSO}_4 \cdot 7\text{H}_2\text{O}$ | 0.181 |
| $\text{CuSO}_4 \cdot 5\text{H}_2\text{O}$ | 0.010 |
| $\text{H}_3\text{BO}_3$ | 0.010 |
| $\text{Na}_2\text{MoO}_4 \cdot 2\text{H}_2\text{O}$ | 0.010 |
| $\text{NiSO}_4 \cdot 6\text{H}_2\text{O}$ | 0.111 |

**Table S7.**
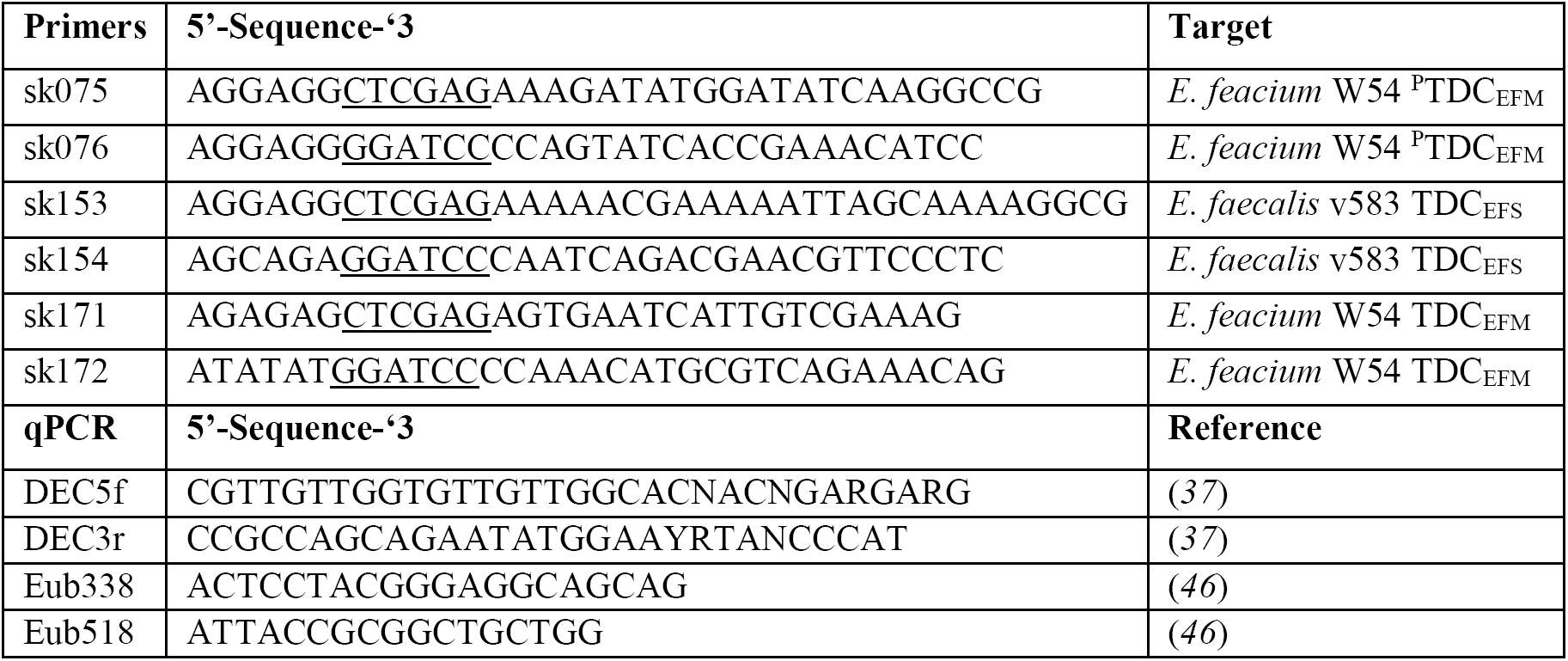
Primer sequences used in this study. Underlined sequences represent restriction sites used.

**Data file S1. Related to Figure 2. Identification of conserved tdcparalogue protein (TDC_EFM_) in all *E. faecium* strains analyzed.** Genome contigs harboring the *tdc* gene cluster of all *E. faecium* strains were extracted from NCBI and aligned using Mauve genome aligner. As comparison, the genome of *E. faecium* W54 is depicted above the alignment results. The paralog TDC from *E faecium* (^P^TDC_EFM_) is shown in orange. Black boxes indicate the TDC gene in all other strains, white bars indicate the single genes, and colored bars indicate conserved gene clusters.

